# Common excluder barley has more than one mechanism to remove Cd from chloroplasts

**DOI:** 10.64898/2026.08.17.745281

**Authors:** Eugene A. Lysenko, Irina F. Seregina, Alexander A. Klaus, Alexander V. Kartashov

## Abstract

Chloroplasts comprise photosynthesis and other important processes. Plants protect chloroplasts from stresses including Cd accumulation. Common terrestrial plants, excluders apply a set of mechanisms to restrict Cd penetration to chloroplasts. Removal of accumulated Cd from chloroplasts should also be a beneficial strategy. However, we do not know whether excluder plant species have ability to remove Cd from chloroplasts. We used barley as a common excluder plant species. To barley plants, we applied a model with two stable isotopes ^111^Cd and ^114^Cd to distinguish Cd accumulated earlier and later. A portion of Cd absorbed by roots continued translocation to shoot for some days after the external source of Cd was changed from one isotope to another. Chloroplasts acquired new portions of Cd and lost part of Cd accumulated earlier; a total Cd content remained rather unchanged. Cd loss from thylakoids was detected *in vivo* and *in vitro*. Cd loss from stroma and envelope was observed *in vivo* but not *in vitro*. Therefore, barley has at least two distinct mechanisms for Cd removal from chloroplasts: one from thylakoids and another from stroma. We hypothesized diverse chlorophagy pathways as a potential mechanism for Cd removal from chloroplasts. Cd accumulation by chloroplasts was mainly light-independent. In chloroplasts, Cd accumulated *in vivo* was tightly bound and mainly located in thylakoids. *In vitro*, chloroplasts from Cd-treated plants accumulated much less Cd than chloroplasts from untreated plants in a previous study. This implies reorganization of transport across chloroplast envelope membranes.

**Highlights:**

- Cd was removed from thylakoids both *in vivo* and *in vitro*
- Cd was removed from stroma and envelope *in vivo* but not *in vitro*
- In chloroplasts, Cd accumulated *in vivo* was tightly bound
- Cd accumulation by chloroplasts was mainly light-independent
- Root barrier slowed down Cd translocation to shoot but not halted it

## 1. Introduction

Distribution of heavy metals is a global problem and Cd is one of the most toxic heavy metals. By origin of contamination, soils can be classified to three types (Baker et al. 2010): 1) extremely rare metalliferous sites of natural origin; 2) mine surroundings that are estimated as 1% of land surface (Prach and Tolvanen 2016); 3) wide areas contaminated by atmospheric and alluvial transfer of industrial emissions and agricultural application of fertilizers and pesticides. Cadmium can be found in metalliferous calamine soils (Wójcik et al. 2017). Many territories are contaminated by Cd of anthropogenic origin (Pan et al. 2010; Zou et al. 2021). Cd is mainly produced as a by-product of Zn; production of one ton of Zn is accompanied by production of 3 kg of Cd (Kabata-Pendias and Mukherjee 2007). The major sources of Cd pollution are atmospheric transfer of industrial emission and phosphate fertilizers (Kabata-Pendias and Mukherjee 2007). The global average Cd concentration in soil is estimated as 0.5 mg kg^-1^ (Kabata-Pendias and Mukherjee 2007) and Cd mobility in soil is rather high (Gaillardet et al. 2003; Kabata-Pendias and Mukherjee 2007; Jigyasu et al. 2020).

Terrestrial plants mostly absorb Cd by roots from soil. The majority of plant species restrict Cd translocation from roots to shoots; they prevent Cd accumulation in aboveground part and called excluders (Baker 1981). Relatively small group of terrestrial plants accumulate the most of Cd in their shoots; these plants are called accumulators (Baker 1981) or hyperaccumulators (Baker and Brooks 1989). Common excluder plants own many mechanisms preventing Cd accumulation in the most important cellular compartments. They bind Cd by cell wall polysaccharides and prevent Cd translocation across plasmalemma; in cytosol, Cd is excluded to vacuole (across tonoplast) and outside of a cell (across plasmalemma) or bound by phytochelatins and metallothioneins (Sanita di Toppi and Gabbrielli 1999; Lin and Aarts 2012). Chloroplasts are a site of energy generation that is required for many protective mechanisms (*e.g.*, synthesis of phytochelatins and metallothioneins). The mentioned mechanisms realize a barrier function preventing Cd penetration to chloroplasts. In leaf mesophyll, central vacuole occupies 73-79% and chloroplasts occupy 16-19% of cell volume (Winter et al. 1993, 1994); therefore, chloroplasts represent the largest site of central metabolism in leaf mesophyll. *In vivo*, chloroplasts of dicots (Baryla et al. 2001) and monocots (Pietrini et al. 2003; Lysenko et al. 2015) accumulated small amount of Cd. Obviously, this is a result of a barrier function restricted Cd uptake by chloroplasts in leaves. However, do plants have an ability to exclude Cd from chloroplasts?

Plants do not have specific Cd transporters. Proteins belonging to families NRAMP, HMA, ZIP, YSL and ABC translocate essential cations Fe, Zn, and Mn across membranes and can also transport Cd and some other heavy metals (Liu et al. 2024). Gene family HMA (Heavy Metal ATPase) is divided to two parts by a similarity to homologous bacterial transporters: one subfamily specific to Cu/Ag and another subfamily specific to Zn/Co/Cd/Pb (Axelsen and Palmgren 2001). Knockdown or knockout of HMA1 gene increased Cd content in chloroplasts of Cd and Zn hyperaccumulator species *Sedum plumbizincicola*; the authors concluded that SpHMA1 perform Cd export from chloroplasts (Zhao et al. 2019). However, HMA1 orthologous group is well-defined in dicots and monocots (Huang et al. 2022; Zang et al. 2023; Shao et al. 2025) and was studied in excluder species. HMA1 is located in chloroplast envelope and mainly participate in transport of Cu and Zn (Seigneurin-Berny et al. 2006; Kim et al. 2009; Mikkelsen et al. 2012) while Cd, Co, Mn, Fe, and Ca can also be transported by HMA1 (see chapter 4.4). The direction of transport remains controversial. Results of one group lookes more trustworthy; according to them, HMA1 transports Cu (but not Zn) in chloroplasts (Seigneurin-Berny et al. 2006; Boutigny et al. 2014). According to others, HMA1 transported Zn (but not Cu) out of chloroplasts under excess of Zn (Kim et al. 2009) and SpHMA1 transported Cd out of chloroplasts under excess of Cd (Zhao et al. 2019).

The result obtained by Zhao with colleagues (2019) is perspective for further studies in Cd hyperaccumulators. However, we have no data on Cd exclusion from chloroplasts in common Cd excluders. Therefore, we carried out a set of model experiments to reveal possible decrease of Cd content in chloroplasts. For some reasons, we selected barley plants. Barley accumulates rather high amount of Cd in chloroplasts *in vivo* (Lysenko et al. 2015). We had performed experiments with Cd uptake by barley chloroplasts *in vitro* and studied Cd distribution between stroma and thylakoids both *in vivo* and *in vitro* (Lysenko et al. 2019). In the current study, we have traced Cd content in chloroplasts after Cd removal from mineral media. Next, we have applied isotopes ^111^Cd and ^114^Cd and verified Cd content in chloroplasts during continuous Cd action. Different isotopes enable to distinguish Cd accumulated initially and later. The latter approach was applied *in vivo*, *in organo* (shoots without roots) and *in organello* (isolated chloroplasts *in vitro*). After Cd accumulation *in vivo* and *in vitro*, ^111^Cd and ^114^Cd were analyzed in thylakoids and in “stroma+envelope” fraction that originated more interesting results. The data obtained is shown and discussed below.

## 2. Materials and Methods

### 2.1. General growth conditions

Barley (*Hordeum vulgare* L. cv. Luch) seedlings were grown in phytotron chambers at 21°C, 200–350 µmol photons m^-2^ s^-1^ and a photoperiod of 16 h light/8 h dark on modified Hoagland medium under continuous aeration (Lysenko et al. 2019). Preliminary, caryopses were kept for 2–3 days at 4°C in darkness on a filter paper moistened with 0.25 mM CaCl_2_. Imbibed caryopses were transferred on plastic mesh that was on top of a vessel containing mineral medium; caryopses were covered with wet filter paper whose edges were immersed in the medium. The caryopses were placed in growth conditions, and plant age was determined from this time. Three days later, the filter paper was removed, and Cd was added to the hydroponic media. The Hoagland medium with monoisotopic Cd salts was reused; Cd content in the medium was verified and demonstrated no substantial changes. After a week of growth, the medium was supplied with extra 3 mM KNO_3_, 2 mM MES, and 1 mM НNO_3_; for cut shoots (see chapter 2.4) additional 3 mM KNO_3_ was added; in a middle of a next week, 1 mM KNO_3_ and 0.5 mM НNO_3_ were added.

### 2.2. Recovery experiment

CdSO_4_ was introduced to the final concentration 80 µM and barley plants were grown until 9^th^ day. Then, the roots were rinsed and plants were transferred to fresh portion of the medium with no Cd addition; plants further grown until 15^th^ day.

Chloroplasts were isolated from the first leaves. The first leaves were quickly detached and kept on ice covered with paper during the collection. All further procedures performed at 4°C in shaded light; all intermediary products were kept on ice. All reusable equipment was presoaked twice in 2 mM EDTA and twice in 0.1 N HCl for at least 2 h each time. The protocol of chloroplast isolation and fractionation described earlier in more details (Lysenko et al. 2019).

Leaves were homogenized in buffer A (50 mM Tris–HCl, pH 8.0, 0.4 M sorbitol, 15 mM NaCl, 2 mM EDTA, 5 mM β-mercaptoethanol) and filtered through one layer of cheesecloth and two layers of miracloth (Calbiochem–Behring, United States). The homogenate was centrifuged for 3 min at 3,500 rpm (Hitachi CR22G III, R7S, Japan) in tubes with flat bottom. Supernatant was discarded and pellet was gently resuspended in buffer A; resuspending was carried out by a gentle agitation of buffer A in tube solely. Percoll (GE Healthcare, United States) was diluted with buffer P (buffer A plus 3% w/v polyethylene glycol 6,000, 0.5% w/v bovine serum albumin, 0.5% w/v Ficoll 400). Chloroplasts were carefully loaded on two-step (20/40%) Percoll gradient and centrifuged for 12 min at 5,800 rpm (Janetzki, K-23, East Germany). Chloroplasts were collected at the interface of 20% and 40% layers and resuspended in 50 ml of buffer A in screw- cap “falcon” tube (Eppendorf, Germany); a sample for chlorophyll (Chl) determination collected. Chloroplasts were sedimented at 5,500 rpm (Eppendorf 5810 R, Germany) for 3 min and the most of supernatant discarded; next, the tubes were centrifuged another 2 min and the rest of supernatant was carefully removed with an automated pipette avoiding loss of chloroplasts. The pellet was kept at -20°C.

### 2.3. Accumulation of Cd isotopes *in vivo*

We used Cd salts enriched with a single isotope: ^111^CdCl_2_ with the isotope enrichment to 95.2% of all Cd isotopes and ^114^CdSO_4_ with the isotope enrichment to 97.06% of all Cd isotopes (FGUP Kombinat Electrochimpribor, Russia). The monoisotopic Cd salts were always introduced with an equimolar amount of magnesium salt to level out anionic content: (^111^CdCl_2_ + MgSO_4_) or (^114^CdSO_4_ + MgCl_2_).

Cd was introduced to the final concentration 80 µM Cd. To verify possible isotope effect, the experiments performed in reciprocal pairs. A) Plants were grown on the medium with 80 µM ^111^Cd from 3^rd^ to 8^th^ day; then, the roots were rinsed, plants were transferred to the medium with 80 µM ^114^Cd and grown until 16^th^ day. B) Plants were grown on the medium with 80 µM ^114^Cd from 3^rd^ to 8^th^ day; then, the roots were rinsed, plants were transferred to the medium with 80 µM ^111^Cd and grown until 16^th^ day.

Chloroplasts were isolated from the first leaves. The procedure was as described in subchapter 2.2. After Percoll gradient, chloroplasts were resuspended in 150 ml of buffer A and centrifuged for 2 min at 3,500 rpm (Hitachi CR22G III, R7S, Japan) in tubes with flat bottom. Supernatant was thoroughly removed using a pipette and pellet was gently resuspended in 8 ml of buffer A; a sample (0.1 ml) for Chl determination collected. The chloroplasts were lysed by hypo-osmotic shock: they were diluted with 80 ml of 10 mM Tris pH 8.0 and stirred gently for 15 min. Then, 0.9 ml of 4 M NaCl was added for better pellet precipitation and the thylakoids were sedimented by sequential centrifugations for 3 min at 5,500 rpm (Eppendorf 5810 R, 3,500 *g*) in 50 ml “falcon” tube. Supernatant was thoroughly collected, transferred to a centrifuge tube, and centrifuged for 3 min at 22,000 rpm (Hitachi, R22A2, Japan) to remove broken thylakoids. Supernatant was transferred to flask, acidified to 1% (HNO_3_ HYPERPUR, PanReac/AppliChem, Canada) and kept at +4°C. Thylakoids kept in “falcon” tube at -20°C.

### 2.4. Accumulation of Cd isotopes “*in organo*”

Plants were grown on the medium with 80 µM Cd from 3^rd^ to 8^th^ day. Then, the shoots were cut at the caryopses level to remove roots; these shoots were placed vertically into layer (1.5-2 cm, 0.75 L) of the medium with 30 µM Cd; the cut shoots were grown until 14^th^ day. The experiments performed in reciprocal pairs. A) Plants were grown on the medium with 80 µM ^111^Cd; then, cut shoots were placed on the medium with 30 µM ^114^Cd. B) Plants were grown on the medium with 80 µM ^114^Cd; then, cut shoots were placed on the medium with 30 µM ^111^Cd.

The stems were cut off; chloroplasts were isolated from the first (major) and second (minor) leaves. The procedure was as described in chapters 2.2 and 2.3 with few modifications to increase chloroplast yield. The leaf homogenate was centrifuged for 5 min at 3,500 rpm (Hitachi). Percoll gradient was centrifuged for 15 min at 5,800 rpm (Janetzki). After Percoll gradient, chloroplasts were resuspended in buffer A and centrifuged for 5 min at 3,500 rpm (Hitachi). The pellet was resuspended in 6 ml of buffer A; a sample (0.1 ml) for Chl determination collected. The chloroplast suspension kept in 15 ml “falcon” tube at -20°C and acidified to 1% (HNO_3_ HYPERPUR) prior to evaporation.

### 2.5. Accumulation of Cd isotopes *in vitro*

Cd was introduced to the final concentration 80 µM and plants were grown until 9^th^ day. Chloroplasts were isolated from the first (major) and second (minor) leaves (stem was cut off) and incubated 1.5 h in a buffer with 25 µM Cd. The experiments performed in reciprocal pairs. A) Plants were grown on the medium with 80 µM ^111^Cd; chloroplasts were incubated in a buffer with 25 µM ^114^Cd. B) Plants were grown on the medium with 80 µM ^114^Cd; chloroplasts were incubated in a buffer with 25 µM ^111^Cd.

Chloroplasts were isolated as described in chapters 2.2 and 2.3 with some modifications. In all buffers, Tric-HCl was replaced with Tricine-NaOH. The leaf homogenate was centrifuged for 3 min at 3,500 rpm (Hitachi). After Percoll gradient, chloroplasts were washed from Percoll and EDTA. Chloroplasts were resuspended in 200 ml of buffer B (50 mM Tricine-NaOH, pH 8.0, 0.4 M sorbitol, 15 mM NaCl) and centrifuged for 5 min at 3,500 rpm (Hitachi) in tubes with flat bottom. Supernatant was thoroughly removed using a pipette. The pellet was washed for 10 min on ice in 50 ml of buffer B with gentle agitation and partial resuspension. Next, it was centrifuged for 1 min at 3,500 rpm (Hitachi) and supernatant was removed using a pipette. Chloroplasts were gently resuspended in buffer B; equal portions of ice-cold chloroplasts (6 ml) were introduced to 39 ml of pre-warmed (21°C) buffer B; Cd concentration was 0 (control) or 25 µM. Chloroplasts were slowly rotated in 50 ml “falcon” tubes on a rotamix for 90 min at 21°C and 70 μmol photons m^-2^ s^-1^. Inside the tubes, light intensity was 50 μmol photons m^-2^ s^-1^; inside tubes covered with black paper, light intensity was 0 μmol photons m^-2^ s^-1^ (LI-250A with PAR detector, LI-COR, USA).

Next, the chloroplasts were cedimented for 1,5 min at 3,500 rpm (Hitachi) in tubes with flat bottom; supernatant was thoroughly removed using a pipette. The pellet was washed with buffer B during 30 min on ice; in one of equal Cd containing variants, 2 mM EDTA was added (EDTA post-washing). The chloroplasts were mostly resuspended; we avoided complete resuspension saving intactness. Chloroplasts were centrifuged for 1,5 min at 3,500 rpm (Hitachi). Supernatant was thoroughly removed using a pipette; the pellet was resuspended in 6 ml of buffer B; a sample (0.1 ml) for Chl determination collected. The chloroplasts were lysed by dilution in 60 ml of 10 mM Tricine-NaOH pH 8.0 and gentle stirring for 15 min. Then, 0.7 ml of 4 M NaCl was added and the procedure was completed as described in chapter 2.3.

### 2.6. Chl measurement

Chlorophylls (Chl a and Chl b) and carotenoids were extracted with 80% acetone, concentrations were measured according to Lichtenthaler and Wellburn (1983). In a leaf blade, the terminal 1-cm segment was removed, and the following 1-cm segment was analyzed.

### 2.7. Measurement of metals

Plant organs were dried at 60°C overnight. Large volume supernatant fractions containing components of stroma and pieces of envelope were evaporated according to Lysenko et al. (2019). Dried organ samples (50–100 mg), pellets of chloroplasts and thylakoids, and concentrated supernatants (stroma + envelope) were transferred to borosilicate tubes (Schott Duran, Germany), mixed with 1.5 mL of 69% nitric acid and 0.6 mL of 70% perchloric acid (HYPERPUR, PanReac/AppliChem, Canada), and incubated overnight. Chloroplasts in 6 ml of buffer A were evaporated with the acids in the borosolicate tubes. The next day, the samples were heated for 1.5 h at 150 °C, then for 2 h at 180 °C; thereafter, 50 µL of concentrated hydrogen peroxide were added, and the samples were incubated overnight. Afterward, the samples were adjusted to the final volume (5 or 10 mL) with deionized water (18.2 МΩ cm^-1,^ Millipore Simplicity, France) acidified with HNO_3_ (1:100).

Concentrations of ^111^Cd and ^114^Cd were measured on a quadrupole ICP-MS spectrometer Agilent 7500c (Agilent Technologies, Japan). The device was controlled by a personal computer using the software ICP-MS-TOP. The detailed parameters and operating conditions of the mass- spectrometer are presented in Suppl. Table S1. ^111^Cd and ^114^Cd standard solutions were prepared from ^111^CdCl_2_ (95.2%) and ^114^CdSO_4_ (97.06%) (FGUP Kombinat Electrochimpribor, Russia) and used for calibration. Deionized water (18.2 МΩ cm^-1,^ Millipore Simplicity, France) used for preparation of all solutions. A single-element 1000 μg mL^-1^ Rh standard solution (High-Purity Standards, USA) used for preparation of the internal standard solutions. In other cases, metals were measured with an atomic absorption spectrophotometer (AA-7000, Shimadzu, Japan) equipped with a hollow-cathode lamps (Hamamatsu, Japan). Prior to the analysis, samples were appropriately diluted with deionized water acidified with HNO_3_ (1:100). For the measurement of Ca and K, the diluting solutions included LaCl₃ (10 g L⁻¹) or LiCl (1 g L⁻¹) respectively to mitigate ionization interference. Parallel sample blank probes were prepared in same manner for background correction.

### 2.8. Equations

Estimation of metal portion in chloroplasts (%) as a part of total accumulation of this metal in leaves performed according to Lysenko et al (2015, 2019) by the equation:

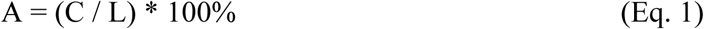

where C is content of this metal in chloroplasts (nmol/mg Chl) and L is content of this metal in leaves (nmol/mg Chl).

Corrections for Chl decrease in chloroplasts were calculated by two ways. Coefficient B shows Cd increase in the first leaves from 8^th^ to 16^th^ day:

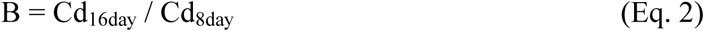

The same quantities of Cd were calculated per a single leaf, DW, FW, or Chl: B_leaf_ ≈ B_DW_ < B_FW_ << B_Chl_.

In leaves, Chl was calculated per FW; therefore, difference between B_FW_ and B_Chl_ reflects an impact of Chl change (Cd values are the same)

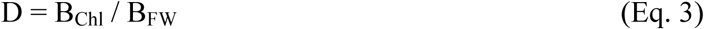

First type correction for Chl decrease was calculated for Cd content in chloroplasts of 16- day-old plants:

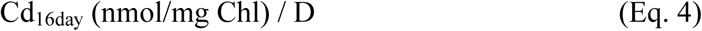

Metal portion in chloroplasts (Eq. 1) should not be sensitive to physiological changes in Chl content because metal contents in both leaves (L) and chloroplasts (C) were normalized to Chl content in same leaves. Coefficient E shows decrease of Cd portion (%) in chloroplasts (Eq. 1) from 8^th^ to 16^th^ day:

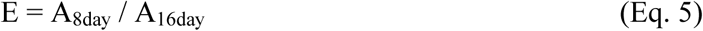

Cd portion (%) in chloroplasts can be decreased by either Cd decrease in chloroplasts (F) or Cd increase in leaves (B).

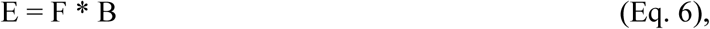

B_leaf_ ≈ B_DW_ shows proper value of Cd increase; B_FW_ was slightly enlarged due to water loss by the leaves. We were not sure that the latter effect can be applicable to chloroplasts; therefore, we used B_FW_ for calculation. If B_leaf_ or B_DW_ will be applied, then F and the correction effect will be slightly larger. Consequently:

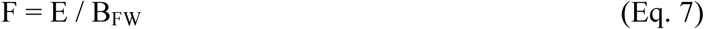

Second type correction for Chl decrease was calculated for Cd content in chloroplasts of 16-day-old plants compared to that in 8-day-old plants:

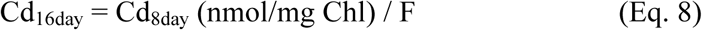

### 2.9. Statistics

Each experiment with Cd isotopes performed in two reciprocal variants (see above). Each recovery experiment and each reciprocal variant carried out in four or five independent repeats. Light dependence of Cd accumulation in chloroplasts *in vitro* studied once in two independent reciprocal variants. The data processed using the Excel (Microsoft) software. The significance of differences between mean values verified with *t*-test.

## 3. Results

### 3.1. Recovery experiment

We began with a simple model. Barley plants were grown from the third to ninth day accumulating Cd; next, they we grown for another six days on the same medium without Cd. A preliminary experiment showed that the first leaves completed their growth to the eighth day and demonstrated no sign of ageing until 16-19^th^ days. Therefore, the first leaves were rather constant during this period.

In the absence of external Cd, the first leaves demonstrated symptoms of recovery. Dry weight (DW) remained unchanged; water content increased; Chl a/b ratio and Chl/carotenoid ratio decreased (Suppl. Table S2). The contents of K, Na, and Ca increased in leaves, K increased in the chloroplasts, while Cu content decreased in both leaves and chloroplasts (Suppl. Table S3). However, no Cd decrease was revealed; instead, insignificant gradual Cd increase observed in both leaves and chloroplasts (Fig. 1, Suppl. Table S3). Possibly, a bulk of Cd accumulated in the roots went on translocation to the leaves despite Cd absence in the mineral medium. To verify this possibility, we used sophisticated experimental model.

**Figure 1.**
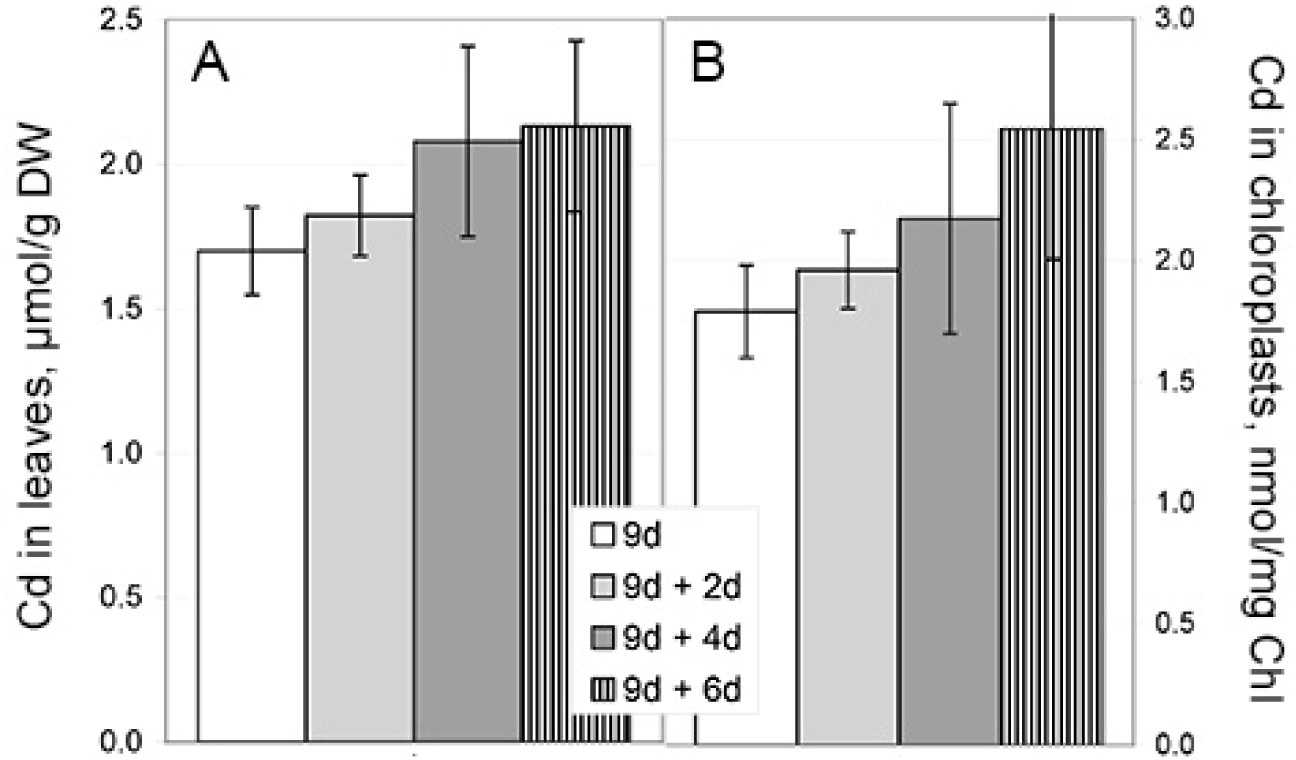
Changes of Cd content at Cd-free mineral media in leaves and chloroplasts. Barley plants were grown at Cd-containing medium from third to ninth day; next, they were transferred to Cd-free medium for another six days. A – Cd content in the first leaves; B –Cd content in chloroplasts isolated from the first leaves. 9-day-old plants at Cd (white) and extra 2 (light grey), 4 (dark grey), or 6 (hatched) days at Cd-free medium. Data are given as means ± SE; the differences were insignificant.

### 3.2. Accumulation of “initial” and “late” Cd in organs

Barley plants were grown from 3^rd^ to 16^th^ day at the same concentration of Cd. However, for the first five days (days 3-8) plants were grown on a one Cd isotope and for the next eight days (days 8-16) they were grown on an alternative Cd isotope. We used ^111^Cd and ^114^Cd isotopes and applied them in reciprocal pairs (^111^Cd/^114^Cd and ^114^Cd/^111^Cd, see chapter 2.3) to verify possible impact of isotope-specificity. In each reciprocal pair, we traced a distribution of initially accumulated isotope (“initial Cd”) and of an isotope that was introduced later (“late Cd”).

From 8^th^ to 16^th^ day, barley plants grew; the roots and shoots enlarged both fresh weight (FW) and DW (Suppl. Table S4). The first leaves mainly remained their masses unchanged, lost water content (Suppl. Table S4) and contents of Chl and carotenoids while Chl a/b and Chl/carotenoids ratios remained unchanged (Suppl. Table S5). The reciprocal pairs of treatment (^111^Cd/^114^Cd and ^114^Cd/^111^Cd) demonstrated indistinguishable results with few exceptions (Suppl.

Tables S4 and S5). Probably, ^111^Cd and ^114^Cd imposed the same effect on plant growth; rare significant differences between the reciprocal variants showed no systemic effect and likely caused by random events. However, these (random) significant differences were reflected with the contents of initial Cd in the first leaves (Fig. 2).

**Figure 2.**
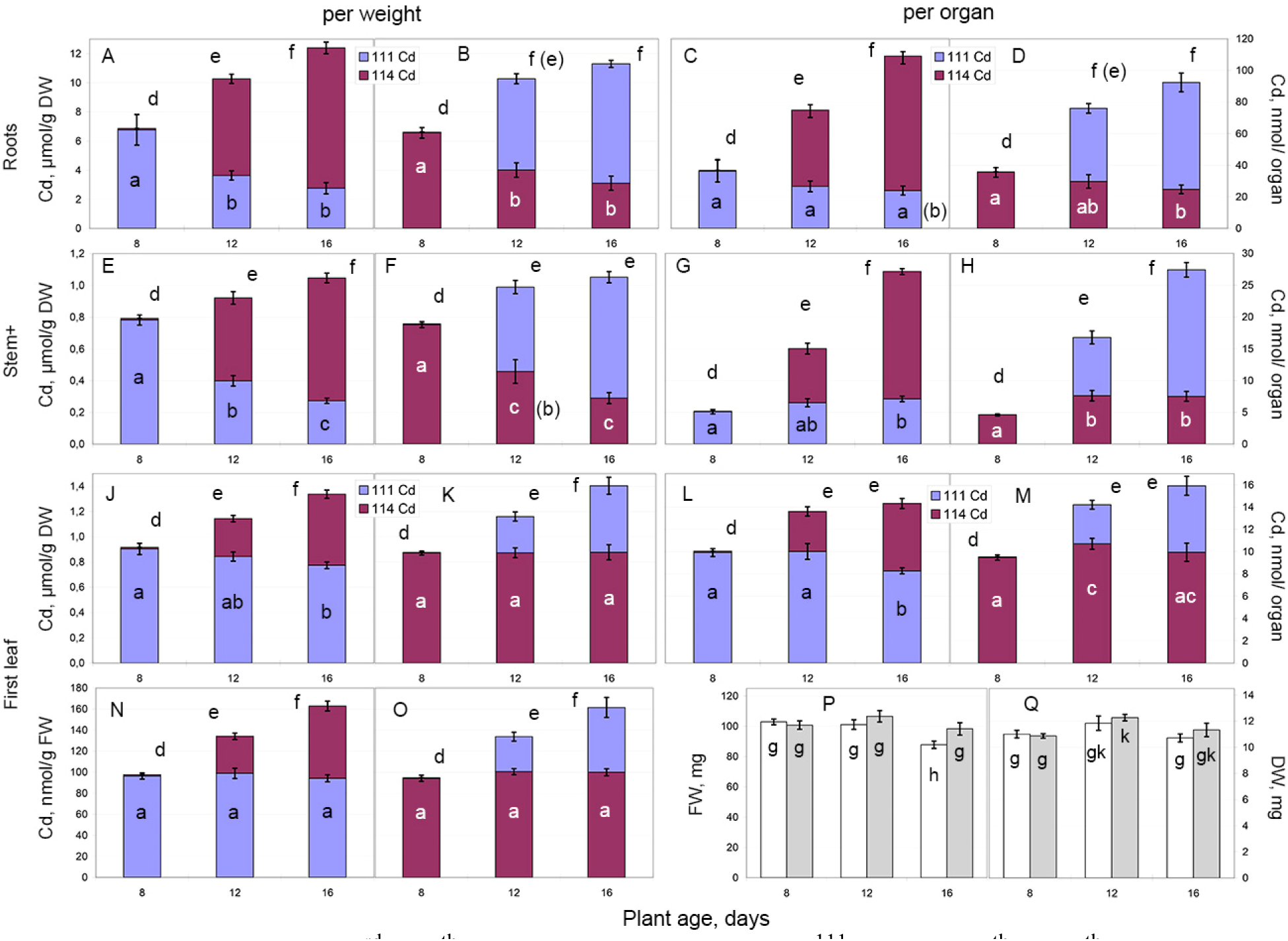
Сontents of ^111^Cd and ^114^Cd in organs of barley plants during reciprocal experiments *in vivo* with “initial” and “late” Cd. A, C, E, G, J, L, N - from 3^rd^ to 8^th^ day, plants were grown at ^111^Cd; from 8^th^ to 16^th^ day, plants were grown at ^114^Cd (experiment ^111^Cd/^114^Cd). B, D, F, H, K, M, O - from 3^rd^ to 8^th^ day, plants were grown at ^114^Cd; from 8^th^ to 16^th^ day, plants were grown at ^111^Cd (experiment ^114^Cd/^111^Cd). A-D – Cd contents in roots (per organ – all roots of a single plant); E-H – Cd contents in stem with leaf sheaths (stem+); J-O – Cd contents in the first leaf (leaf blade). A-B, E-F, J-K - Cd contents per DW; C-D, G-H, L-M - Cd contents per organ; N-O - Cd contents per FW. Blue - ^111^Cd; burgundy - ^114^Cd. Significant differences of Cd contents between variants, p < 0.05: a-c – differences of initial Cd (^111^Cd or ^114^Cd given from 3^rd^ to 8^th^ day); d-f - differences of total Cd (^111^Cd + ^114^Cd); in parenthesis – difference of a datapoint in combined variant (all initial Cd/ all late Cd, N = 10). The contents of late Cd (^111^Cd or ^114^Cd given from 8^th^ to 16^th^ day) always differed significantly at p < 0.05, therefore, remained unmarked (to ease perception). P – FW of the first leaf (blade); Q – DW of the first leaf (blade). Reciprocal experiments: white – ^111^Cd/ ^114^Cd; grey – ^114^Cd/ ^111^Cd. g-k - significant differences between FWs or DWs, p < 0.05. Data are given as means ± SE.

Dynamics of essential metals in barley organs were undistinguishable in the reciprocal pairs of treatment (^111^Cd/^114^Cd and ^114^Cd/^111^Cd); no significant difference observed between reciprocal variants (Suppl. Table S6). Growing organs (roots and stems) increased contents of all metals. Due to the growth, contents of Cu, Mn, Mg, and Na decreased per weight and increased per the whole organ; contents of Zn, Fe, Ca, and K increased per both weight and organ, while the increase per weight was smaller than the increase per organ. The first leaves were rather constant and metal contents changed similarly per both weight and organ. In the first leaves, contents of the most metals increased, K increased in the middle of treatment (12^th^ day), Mg unchanged and Cu decreased (Suppl. Table S6).

From 8^th^ to 16^th^ day, barley accumulated more Cd in all organs; this was achieved due to the uptake of late Cd (Fig. 2). The contents of initial Cd clearly decreased per DW in roots and stems (Fig. 2A-B, E-F); however, this was caused by the growth of these organs. The whole roots of a single barley plant lost 11.6 ± 3.9 nmol of initial Cd (Fig. 2C-D); the roots contained 35.9 ± 3.6 nmol of initial Cd at 8^th^ day and 24.3 ± 1.8 nmol of initial Cd at 16^th^ day (hereinafter combined data of all reciprocal experiments are shown). Stem acquired 2.5 ± 0.5 nmol of initial Cd. Barley stem with leaf sheaths contained 4.8 ± 0.2 nmol of initial Cd at 8^th^ day and 7.3 ± 0.4 nmol of initial Cd at 16^th^ day (Fig. 2G-H). This showed that even if Cd removed from media it is still translocating to the aboveground part from roots.

In the first leaves, initial Cd mainly remained at the same level (∼10 nmol/leaf) though it demonstrated surprising fluctuations (Fig. 2J-O). The content of initial Cd per FW remained constant from 8^th^ to 16^th^ days in both reciprocal pairs of experiment (Fig. 2N-O). In the experiment “initial ^111^Cd/ late ^114^Cd” at 16^th^ day, we observed the decrease of initial Cd content per DW and per organ (Fig. 2J, L) and of the first leaf FW (Fig. 2P). In the reciprocal experiment “initial ^114^Cd/ late ^111^Cd” at 12^th^ day, we observed the increase of initial Cd content per organ (Fig. 2M) and of the first leaf DW (Fig. 2Q). Probably, two opposed processes balance the level of initial Cd in the first leaf: a) translocation of initial Cd from roots to the first leaf and b) translocation of initial Cd from the first leaf to growing leaves. Usually, both translocations are equal and Cd content remains unchanged. When some process prevails, we can observe the decrease of FW and initial Cd at 16^th^ day (experiment ^111^Cd/ ^114^Cd) or the increase of DW and initial Cd at 12^th^ day (experiment ^114^Cd/ ^111^Cd).

### 3.3. Accumulation of “initial” and “late” Cd in chloroplasts *in vivo*

We traced changes of metals in barley chloroplasts from 8^th^ to 16^th^ day in the process of *in vivo* experiment with the initial and late Cd. Chloroplasts were isolated from the first leaves. In stroma, content of bivalent cations remained unchanged, while K level increased drastically (Suppl. Table S7). K content in thylakoids is very low (Lysenko et al. 2019) and required more chloroplasts for proper measurement; K content in chloroplasts is mainly determined by its stromal content. Bivalent cations except Ca are mainly accumulated in thylakoids (Lysenko et al. 2019). In thylakoids and chloroplasts, Cu and Mn contents decreased, Mg and Fe contents unchanged, while Cd demonstrated small significant increase (Suppl. Table S7). Cd increase in the first leaves was larger (Fig. 2) than Cd increase in the chloroplasts, therefore portion of chloroplastic Cd in the first leaves reduced from 2% to 1% (Suppl. Table S7). Chloroplastic portions of other bivalent cations reduced also; at least, chloroplastic portion of Fe decreased for the same reason.

The content of initial Cd decreased in both thylakoids and stroma (Fig. 3). In thylakoids, the loss of initial Cd was smaller than acquisition of late Cd; therefore, the total Cd content increased (Fig. 3A-C). In stroma, contents of lost initial Cd and acquired late Cd were equal; thus, the total Cd content unchanged (Fig. 3D-F). This data showed that the portion of Cd was removed from chloroplasts; however, this portion was rather small. Probably, a portion of initial Cd was translocated from roots and penetrated to chloroplasts that masked some portion of removed initial Cd.

**Figure 3.**
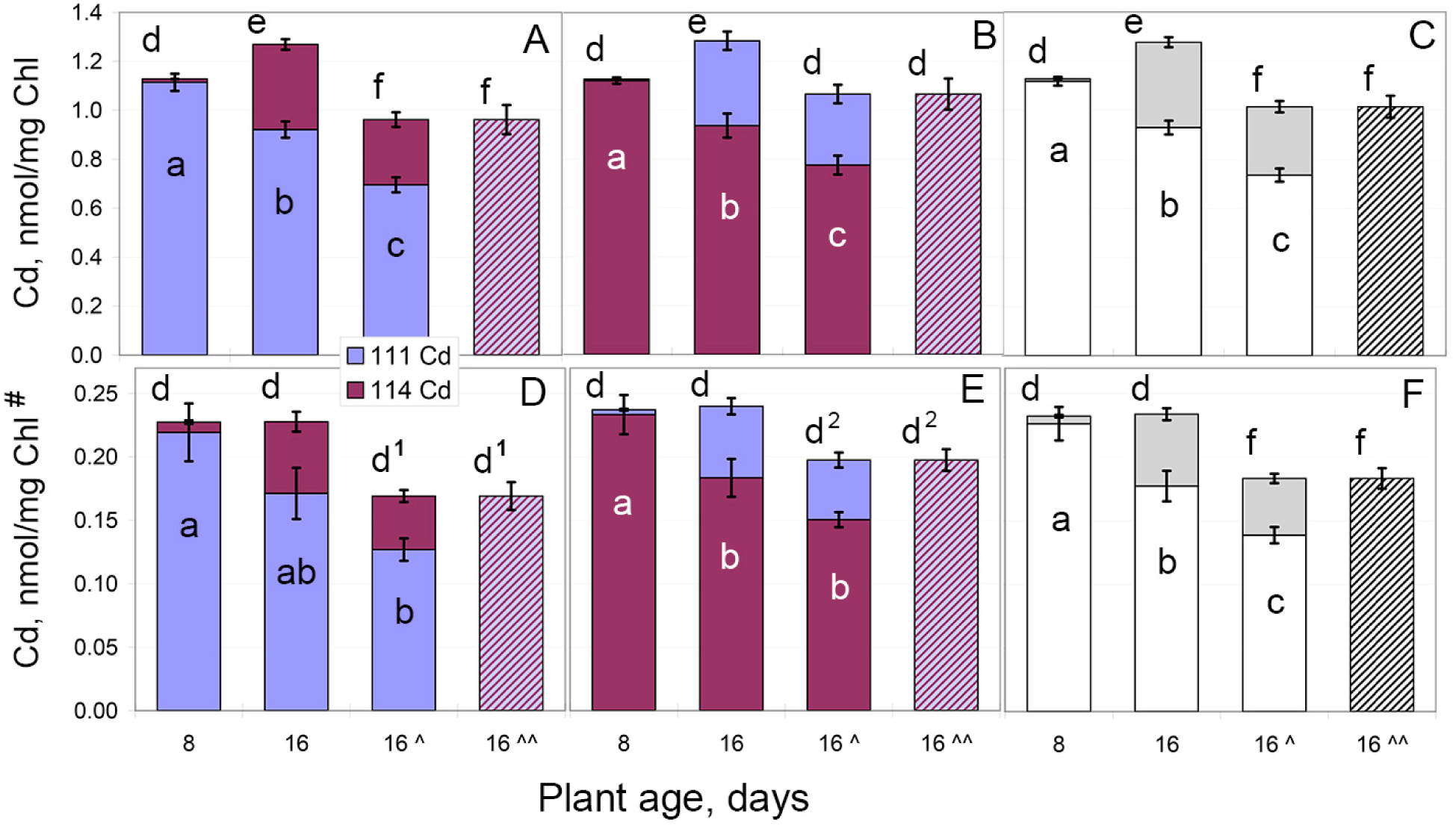
Contents of ^111^Cd and ^114^Cd in barley chloroplasts during reciprocal experiments *in vivo* with “initial” and “late” Cd. A-C – Cd contents in thylakoids; D-F – Cd contents in stroma and envelope. A, D – experiment ^111^Cd/^114^Cd, N = 5; B, E - experiment ^114^Cd/^111^Cd, N = 5; blue - ^111^Cd, burgundy - ^114^Cd, hatched – total Cd (^111^Cd + ^114^Cd). C, F - combined data: white – all initial Cd, grey - all late Cd, hatched – total Cd, N = 10. # - per mg Chl in chloroplasts/thylakoids from which the stromal fraction was separated; ^ - first type correction for Chl decrease (Eq. 4); ^^ - second type correction for Chl decrease (Eq. 8). 1 – difference significant at p = 0.054; 2 – difference significant at p = 0.052. Other designations are the same as in Fig. 2.

Also, Chl content in 16-day-old plants decreased (Suppl. Table S5) that increased the ratio nmol Cd/mg Chl. From 8^th^ to 16^th^ day, Cd content in the first leaves increased by 53-55% per DW and per leaf, by 69% per FW (Fig. 2J-O), and by 114% per mg Chl (not shown). We calculated corrections to compensate the impact of Chl decrease on the values nmol Cd/mg Chl in chloroplasts of 16-day-old plants. The first type correction (Eq. 4) was applied to any Cd isotope. The second type correction (Eq. 8) was applied to total Cd content (^111^Cd+^114^Cd) only, because chloroplasts from 8- and 16-day-old plants have distinct isotope composition. According to corrected values, total Cd content slightly decreased in thylakoids, stroma and chloroplasts of 16-day-old plants because the decrease of initial Cd appeared larger than the acquisition of late Cd (Fig. 3, Suppl. Table S7).

### 3.4. Accumulation of “initial” and “late” Cd in chloroplasts *in organo*

Next, we detached roots before giving to shoots a late Cd. For the first five days (days 3- 8), intact barley plants were grown on a one Cd isotope (80 μM). Then, the shoots were cut and placed on an alternative Cd isotope (30 μM, Fig. 4A). The cut shoots were exposed to late Cd for six days (days 8-14). After one or two days of the exposition, the cut leaves demonstrated visible symptoms of damage and/or ageing, while analogous cut shoots of control plants at Cd-free medium looked healthy for eight and more days. The symptoms were increasing; therefore, we reduced exposition to late Cd from 8 to 6 days. Сhloroplasts were isolated from the first (major) and second (minor) leaves.

**Figure 4.**
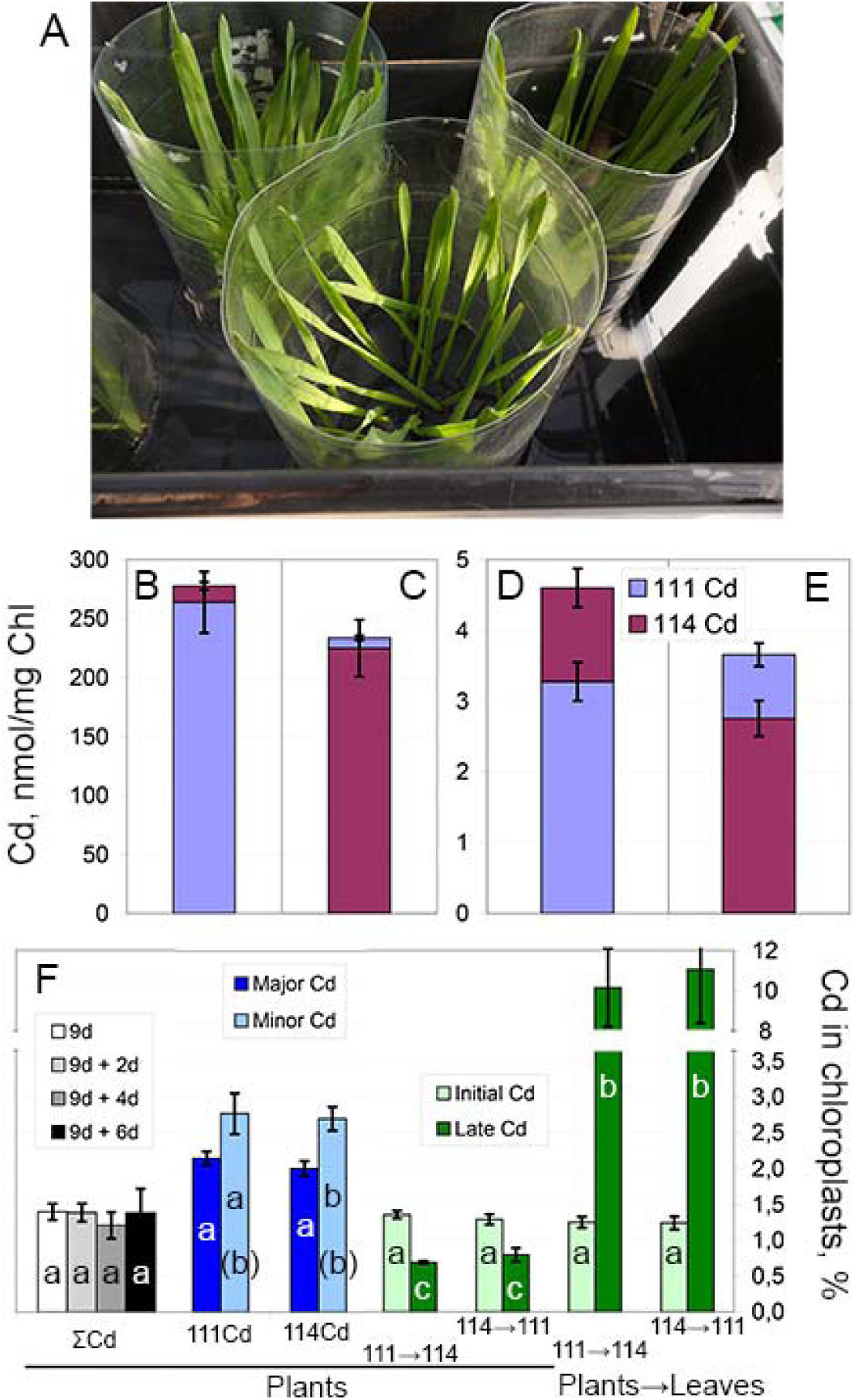
Contents of ^111^Cd and ^114^Cd in barley leaves and chloroplasts during reciprocal experiments *in organo* with “initial” and “late” Cd Barley plants were grown at initial Cd (^111^Cd or ^114^Cd, 80 μM) for five days (days 3-8); next, the shoots were cut and placed on late Cd isotope (^114^Cd or ^111^Cd, 30 μM) for six days (days 8-14). A – general view of cut shoots. B-C – Cd contents in the first leaves; D-E – Cd contents in chloroplasts from the first and second leaves. B, D - experiment ^111^Cd/^114^Cd; C, E - experiment ^114^Cd/^111^Cd. Blue - ^111^Cd, burgundy - 114_Cd._ F – portion (%) of Cd in the chloroplasts comparing with total Cd content in the first leaves; all abovementioned experiments. Plants – experiments with intact plants. ΣCd – general Cd in recovery experiment: 9-day-old plants at Cd (white) and extra 2 (light grey), 4 (dark grey), or 6 (black) days without Cd; a – no significant difference, p < 0.05. 111Cd and 114Cd – 9-day-old plants on the monoisotopic Cd salts with the isotope enrichment 95% and 97% correspondingly; dark blue – distribution of the corresponding major isotope (^111^Cd or ^114^Cd), light blue – distribution of the corresponding second isotope (^111^Cd or ^114^Cd) as one of the minor isotopes; a-b – significant difference between distributions of major and minor isotopes, N = 5, p < 0.05; in parenthesis – significant difference between combined data: all major and all minor isotopes, N = 10, p < 0.05. 111→114 and 114→111 – 16-day-old plants after reciprocal experiments with initial and late Cd: ^111^Cd/^114^Cd and ^114^Cd/^111^Cd correspondingly; light green – distribution of the corresponding initial Cd, dark green – distribution of the corresponding late Cd; a and c – significant difference between distributions of initial and late Cd, p < 0.05. Plants→Leaves – plants were grown on initial Cd (days 3-8) and cut shoots were placed on late Cd (days 8-14); 111→114 and 114→111 – reciprocal experiments with initial and late Cd: ^111^Cd/^114^Cd and ^114^Cd/^111^Cd correspondingly; light green – distribution of the corresponding initial Cd, dark green – distribution of the corresponding late Cd; a-b – significant difference between distributions of initial ant late Cd. Data are given as means ± SE.

In the first leaves of 14-day-old cut shoots, metal contents (Suppl. Table S8) were similar to those in two previous experiments. The contents of most metals resembled those in 16-day-old plants under continuous Cd action (Fig. 2, Suppl. Table S6) while Fe content was similar to that in recovery experiment (Suppl. Table S3). In the first leaves, the bulk of Cd represented by initial Cd (Fig. 4B-C); cut shoots acquired very small amount of late Cd. In the chloroplasts, the contents of Cu, Mn, Mg, and K per mg Chl (Suppl. Table S8) were similar to those in previous experiments (Suppl. Tables S3 and S7); however, proportion (%) of total leaf Cu, Mn, Mg, and K found in chloroplasts (Suppl. Table S8) substantially decreased comparing with previous experiments (Suppl. Tables S3 and S7). Probably, in degrading leaves of cut shoots, the chloroplasts lost both Chl and the metals; thus, the ratio metal/Chl did not changed while contents of these metals in chloroplasts decreased comparing with their whole leaf contents. Using this experimental model, we cannot judge whether initial Cd content in chloroplasts changed or not. This experimental model occurred rather unsuccessful; however, this model clearly showed one fact.

The first leaves of cut shoots acquired small amount of late Cd (Fig. 4B-C). Despite this, late Cd content in their chloroplasts was rather large (Fig. 4D-E) and similar to that accumulated by intact plants (Fig. 3). In barley plants, chloroplasts accumulated 1-2% of all Cd in the first leaves (Fig. 4F; Lysenko et al. 2015, 2019). While in damaged leaves of cut shoots, 10.6 ± 1.5% of late Cd penetrated to chloroplasts (Fig. 4F).

### 3.5. Accumulation of “initial” and “late” Cd in chloroplasts *in vitro*

We studied Cd accumulation by isolated chloroplasts *in vitro*. Barley plants were grown until ninth day on a one Cd isotope (80 μM); then, chloroplasts were isolated from the first and second leaves and incubated 1.5 h in a buffer with 25 µM on an alternative Cd isotope. To our surprise, chloroplasts from Cd-treated plants accumulated much smaller amount of Cd (Fig. 5) than chloroplasts from untreated plants (Lysenko et al. 2019); the difference was 1-2 order of magnitude. Also, the fate of initial Cd was different in the thylakoids and stroma. *In vitro*, thylakoids 1) accumulated major portion of late Cd that was equal to portion of initial Cd accumulated *in vivo* and 2) lost a half of initial Cd (Fig. 5A-B). *In vitro*, stroma accumulated twofold smaller amount of late Cd comparing with thylakoids; the content of initial Cd was low and remained unchanged (Fig. 5C-D).

**Figure 5.**
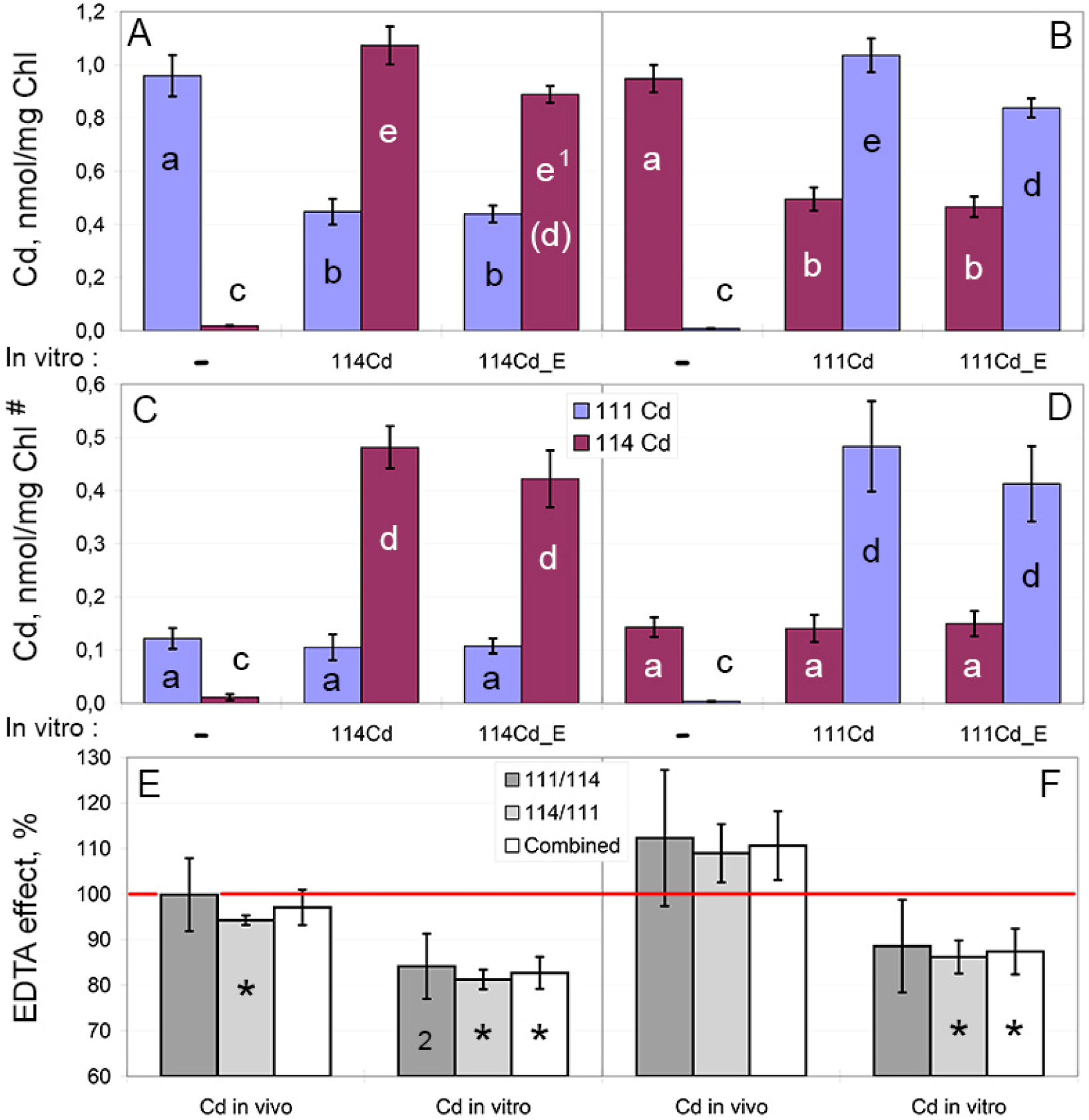
Contents of ^111^Cd and ^114^Cd in barley chloroplasts during reciprocal experiments *in vitro* Barley plants were grown until ninth day on initial Cd (^111^Cd or ^114^Cd, 80 μM); then, chloroplasts were isolated and incubated for 1.5 h in a buffer with late Cd (^114^Cd or ^111^Cd, 25 μM). A-B – Cd contents in thylakoids; C-D – Cd contents in stroma+envelope fractions. A, C – experiment ^111^Cd *in vivo*/ ^114^Cd *in vitro*; B, D – experiment ^114^Cd *in vivo*/ ^111^Cd *in vitro*. Blue - ^111^Cd, burgundy - ^114^Cd. # - per mg Chl in chloroplasts/thylakoids from which the stromal fraction was separated. *In vitro* variants: “-“ – incubation with no Cd addition (control), “Cd” – incubation with the corresponding Cd isotope, “Cd_E” - incubation with the corresponding Cd isotope and post-washing with EDTA. Significant differences of Cd contents between *in vitro* variants, p < 0.05: a-b – differences of initial Cd, c-e – differences of late Cd, in parenthesis – difference between combined data: all major and all minor isotopes, N = 8. 1 – difference significant at p = 0.056. E-F – EDTA post-washing effect on Cd contents. In each repeat of the experiment (A-D), data from the variant “Cd” accepted as 100% and the corresponding data from the variant “Cd_E” calculated as portion (%) of that 100% and showed. Red line marks 100% (Cd content after washing without EDTA). E – thylakoids (from A-B), F – stroma+envelope (from C-D). Dark grey – experiment ^111^Cd *in vivo*/ ^114^Cd *in vitro*; light grey – experiment ^114^Cd *in vivo*/ ^111^Cd *in vitro*; white – combined data: initial Cd *in vivo*/ late Cd *in vitro* (N = 8). “Cd in vivo” – initial Cd accumulated *in vivo*; “Cd in vitro” – late Cd accumulated *in vitro*. * - significant effect of EDTA post-washing, p < 0.05. 2 – difference significant at p = 0.057. Data are given as means ± SE.

We introduced EDTA in the buffer for chloroplast washing after *in vitro* accumulation (post-washing) to test level of loosely bound Cd in the chloroplasts (Lysenko et al. 2019). EDTA post-washing did not change content of initial Cd while removed small portion of late Cd; the latter effect was similar in both thylakoids and stroma but was insignificant in stroma (Fig. 5A- D). We recalculated these data. After post-washing with no EDTA, Cd content was accepted as 100% and the effect of EDTA post-washing was calculated in each repeat of the experiment (Fig. 5E-F). The thylakoids lost 17.3 ± 3.5% of late Cd (Fig. 5E), stroma lost 12.6 ± 5.0% of late Cd (Fig. 5F). The content of initial Cd did not decrease in stroma and in one reciprocal experiment (^111^Cd/^114^Cd) in thylakoids. In another reciprocal experiment (^114^Cd/^111^Cd), initial Cd decreased in thylakoids significantly by ∼5% while the decrease in combined data of both reciprocal experiments was insignificant (Fig. 5E).

Nor Cd, neither EDTA influenced Cu and Mg contents in thylakoids and stroma *in vitro*; data of Fe measurements varied much with no significant changes (Fig. 6A, Suppl. Table S9). Cd treatment *in vitro* decreased K content in stroma (Fig. 6C, Suppl. Table S9). EDTA post-washing caused translocation of a protion of Mn (14%) from thylakoids to stroma (Fig. 6B, Suppl. Table S9). Similar data were obtained in previous experiment (Lysenko et al. 2019) and remained unpublished; we have shown these data in Suppl. Table S10.

**Figure 6.**
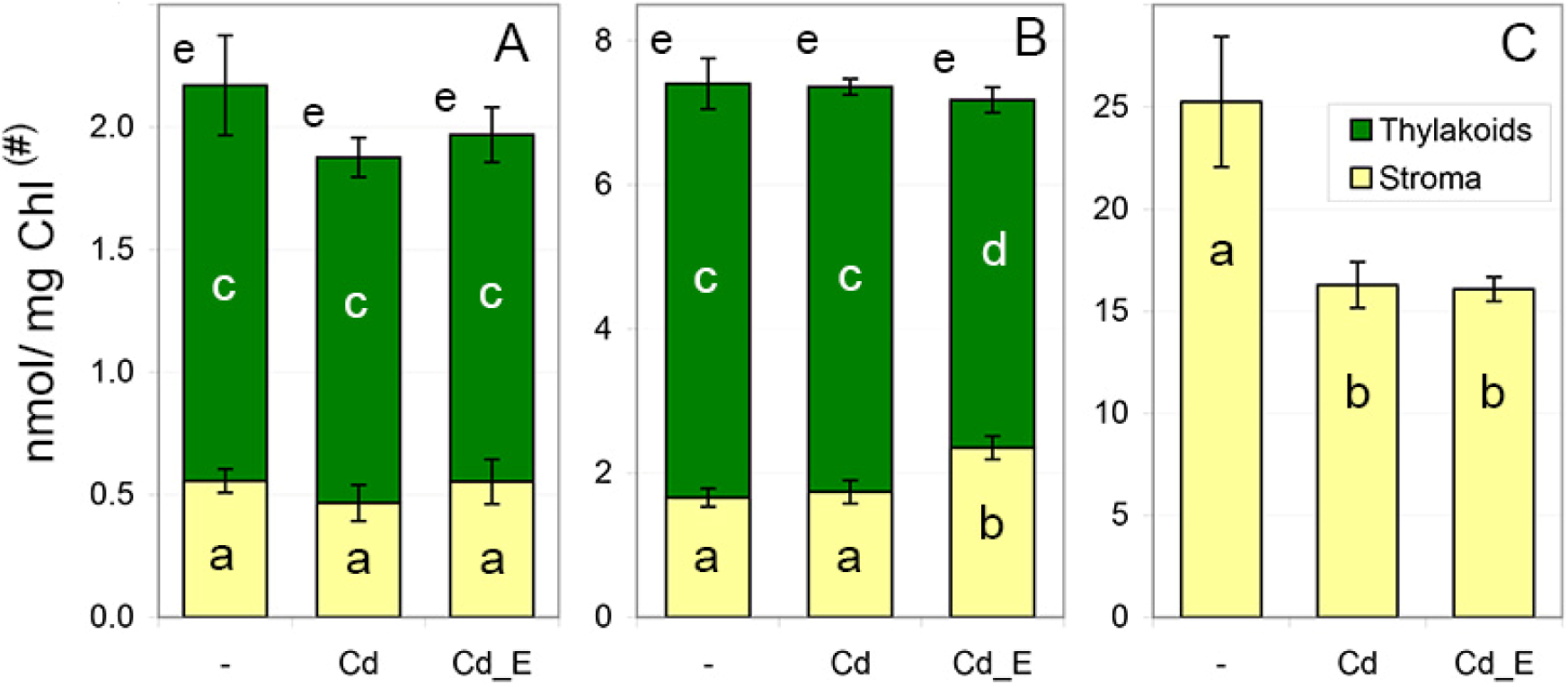
Contents of some essential metals in barley chloroplasts during Cd accumulation *in vitro*The experiment was the same as in Fig. 5. The combined data of reciprocal experiments are shown as means ± SE. The complete information is shown in Suppl. Table S9. A – copper, B – manganese, C – potassium. Green – thylakoids, yellow – stroma+envelope. # and *in vitro* variants designated in Fig. 5 legend. Significant differences of metal contents between *in vitro* variants, p < 0.05: a-b – in stromal fraction, c-d – in thylakoids, e – in whole chloroplasts (no difference). K content in thylakoids is tiny. K contents in stroma is shown for experiments 2-4; for the complete information see Suppl. Table S9.

### 3.6. Light effect on Cd accumulation by chloroplasts *in vitro*

A light effect on Cd accumulation *in vitro* studied in a single experiment. However, results of reciprocal experiments occurred similar; therefore, we showed them. Under illumination and in darkened tubes, barley chloroplast accumulated similar amount of Cd. Stromal Cd content varied to some extent; so, the data were indistinguishably similar (Fig.7D-F). In thylakoids, data were of utmost similarity; therefore, we observed small significant differences. In darkness, thylakoids accumulated less late Cd, lost less initial Cd, and kept same total Cd content comparing with illumination conditions (Fig. 7A-C). Thus, Cd accumulation by chloroplasts mostly is light-independent process, while light can stimulate it to some extent.

**Figure 7.**
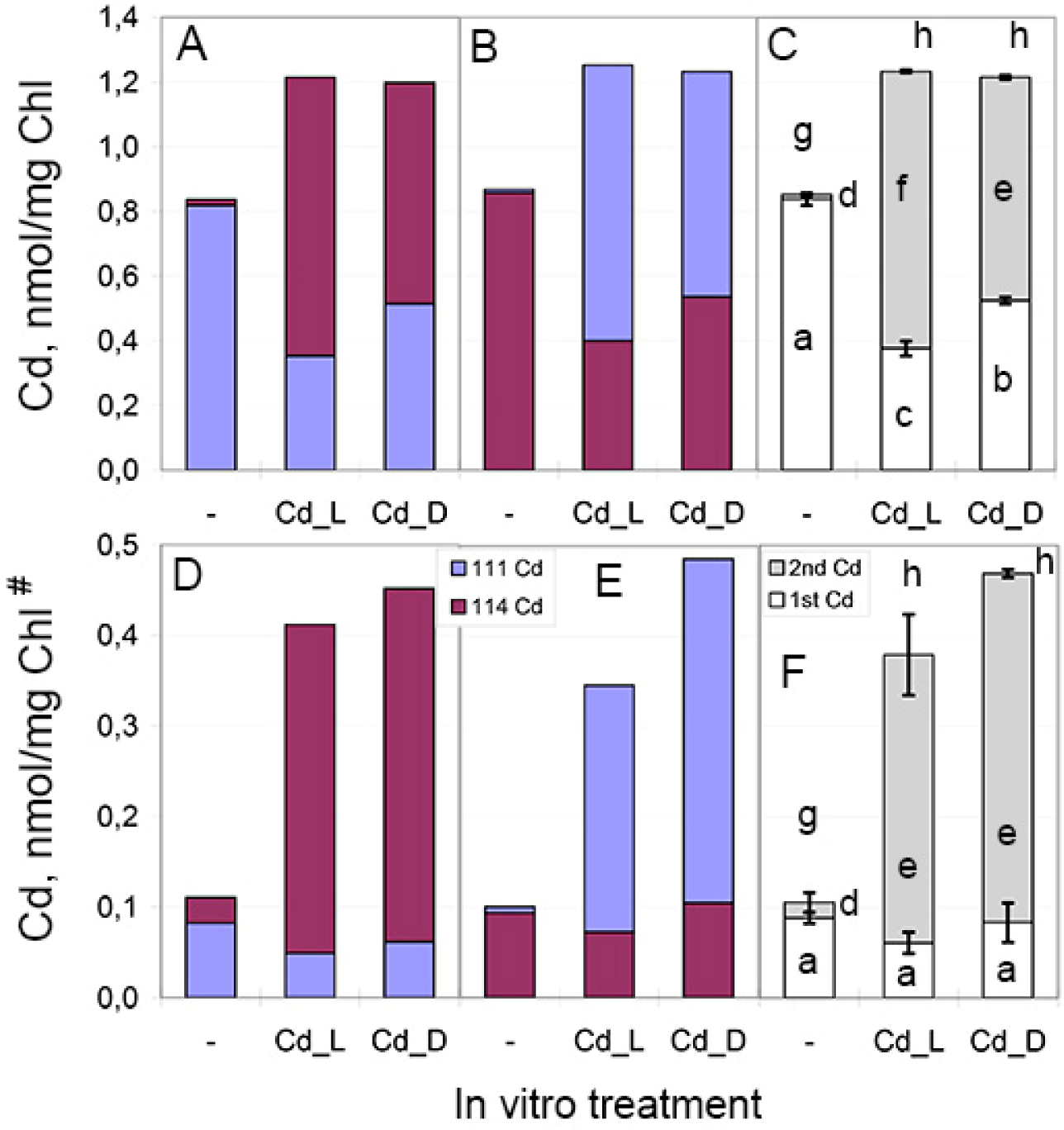
Light dependence of ^111^Cd and ^114^Cd contents in barley chloroplasts in reciprocal experiments *in vitro* The experiment was similar to Fig. 5 while Cd was accumulated *in vitro* in the presence or absence of light. A-C – Cd contents in thylakoids, D-F – Cd contents in stroma+envelope. A, D – single experiment ^111^Cd *in vivo*/ ^114^Cd *in vitro*; B, E – single experiment ^114^Cd *in vivo*/ ^111^Cd *in vitro*; C, F – combined data of experiments: initial Cd *in vivo*/ late Cd *in vitro* (N = 2, means ± SE). Blue - ^111^Cd, burgundy - ^114^Cd, white – initial (1^st^) Cd, grey – late (2^nd^) Cd. # - designated as in Figs 3 and 5. *In vitro* variants: “-“ – incubation with no Cd addition, “Cd_L” – incubation with the corresponding Cd isotope under illumination, “Cd_D” - incubation with the corresponding Cd isotope in darkness. Significant differences of Cd contents between combined *in vitro* variants, p < 0.05: a-c – differences of initial Cd, d-f – differences of late Cd, g-h – differences of total Cd contents (^111^Cd + ^114^Cd).

## 4. Discussion

### 4.1 Cd transport in plants

Barley plants accumulated bulk of Cd in their roots (Fig. 2). When an external source of Cd was eliminated, Cd from roots probably continued translocation to leaves. Data from Fig. 1 assumed this and data from Fig. 2 showed it. During the first five days, roots of a single barley plant accumulated 35.9 ± 3.6 nmol Cd (Fig. 2C-D). For the next eight days, these roots lost 11.6 ± 3.9 nmol of initially accumulated Cd. Of this 11.6 nmol, 2.5 ± 0.5 nmol was translocated to stem. Throughout the second interval of growth, plants acquired late Cd isotope from the medium; however, the content of initial Cd isotope was increased in the stem from 4.8 ± 0.2 nmol to 7.3 ± 0.4 nmol (Fig. 2G-H). Isotope-enriched salts contained other isotopes at low level. For example, ^114^CdSO_4_ contained 3% of all other isotopes; therefore, the content of ^111^Cd was less than 0.5%. If we assume accumulation of minor Cd isotope from solution, we should observe an increase of this isotope content in roots. Instead, we observed decrease of such isotope in roots (Fig. 2C-D). Thus, we have the only explanation. Plants acquired bulk of initial Cd isotope (∼36 nmol) during the first five days; during the next eight days, plants translocated part (∼2.5 nmol) of this bulk to the stem. The rest of “lost” initial Cd (∼9 nmol) was probably transported to leaves; its release to mineral medium is less probable. For example, barley roots lost a portion of ^54^Mn in 5 h while researchers failed to measure its radioactivity in external solution (Pedas et al. 2005). Consequently, lost ^54^Mn was transported to shoot tissues.

The first leaf accumulated near 10 nmol of initial Cd (8d ^111^Cd/^114^Cd: 9.9 ± 0.35 nmol; 12d ^111^Cd/^114^Cd 10.0 ± 0.7 nmol; 16d ^114^Cd/^111^Cd 9.9 ± 0.8 nmol) and 6.0 ± 0.45 nmol of late Cd (Fig. 2L-M). Likely, rather constant ∼10 nmol level resulted from dynamic balance of two processes. We suppose that initial Cd was simultaneously transported from roots to the first leaves and from the first leaves to growing leaves. When the influx and outflow were unequal then the level changed. On 12^th^ day in the experiment ^114^Cd/^111^Cd solely, the first leaves increased DW but not water content and FW (Fig. 2P-Q, Suppl. Table S4); the content of initial ^114^Cd remained unchanged per DW (Fig. 2K) and increased per a leaf (Fig. 2M); the increase per FW was insignificant (Fig. 2O). In this case, transport into the first leaves was probably larger than drain out of them. On 16^th^ day in the experiment ^111^Cd/^114^Cd solely, the first leaves lost water content and FW, while DW remained unchanged (Fig. 2P-Q, Suppl. Table S4); the content of initial ^111^Cd remained unchanged per FW (Fig. 2N) and decreased per DW (Fig. 2J) and per a leaf (Fig. 2L). In this case, transport into the first leaves was probably smaller than drain out of them in this case. Likely, both effects were caused by random events; however, they enable us to see imbalance of transport into and out of the first leaves.

In summary, when we removed external source of initial Cd, it was still transported from roots to stem and leaves. Therefore, “root barrier” does not halt Cd translocation to shoot but slow down this process.

### 4.2 Cd accumulation by chloroplasts

*In vivo*, Cd mostly situated in thylakoids. Of all Cd in chloroplasts, thylakoids contained 80% in 9-day-old (Lysenko et al. 2019), 83% in 8-day-old, and 84.6% in 16-day-old barley plants (Suppl. Table S7). *In vitro*, the portion of Cd penetrated to thylakoids was twice as large compared to Cd portion penetrated to stroma (Fig. 5, Suppl. Table S9). This is in line with the distribution of other bivalent cations inside barley chloroplasts. The largest portions of Cu, Mn, Fe, Mg, and Zn were also located in thylakoids (Suppl. Table S7, Lysenko et al. 2019). Solely, Ca in untreated and Cu- and Fe-treated barley plants was mainly located in stroma, however, in Cd-treated plants, Cd was mainly situated in thylakoids. To the contrary, monovalent cation K was mostly located in stroma and envelope (Lysenko et al. 2019); therefore, we failed to measure it in thylakoids in the current study (Suppl. Tables S7 and S9).

We were unable to replace the most of initial Cd by late Cd. From 8^th^ to 16^th^ day, Cd content in the first leaves increased by about 60% due to accumulation of late Cd and rather constant level of initial Cd (Fig. 2J-O). On 16^th^ day, late Cd constituted 40 ± 2% of all Cd in the first leaves. Limited addition of late Cd to leaves caused limited addition of late Cd to chloroplasts. On 16^th^ day, late Cd constituted 24.3 ± 1.6% of all Cd in stroma and envelope and 27.2 ± 1.5% of all Cd in thylakoids (Fig. 3). In cut shoots, late Cd constituted 4.4 ± 0.7% (Fig. 4B-C); in their chloroplasts, late Cd constituted 26.0 ± 2.3% (Fig. 4D-E). In damaged leaves of 14-day-old cut shoots, 10.6 ± 1.5% of late Cd was observed in chloroplasts; whereas barley chloroplasts accumulated 1-2% of all Cd in the first leaves usually (Fig. 4F). Apparently, the damaged leaves exhausted ability to prevent Cd penetration to chloroplasts and/or remove Cd from chloroplasts. It underlies importance of these mechanisms.

Chloroplasts from Cd-treated plants demonstrated a resistance to Cd penetration. *In vitro*, chloroplasts from untreated plants accumulated large (at 5 μM Cd) or huge (at 100 μM Cd) amount of Cd (Lysenko et al. 2019); however, chloroplasts from Cd-treated plants accumulated small amount of Cd at 25 μM Cd (Fig. 5, Suppl. Table S9). These data are summarized in Fig.8.

**Figure 8.**
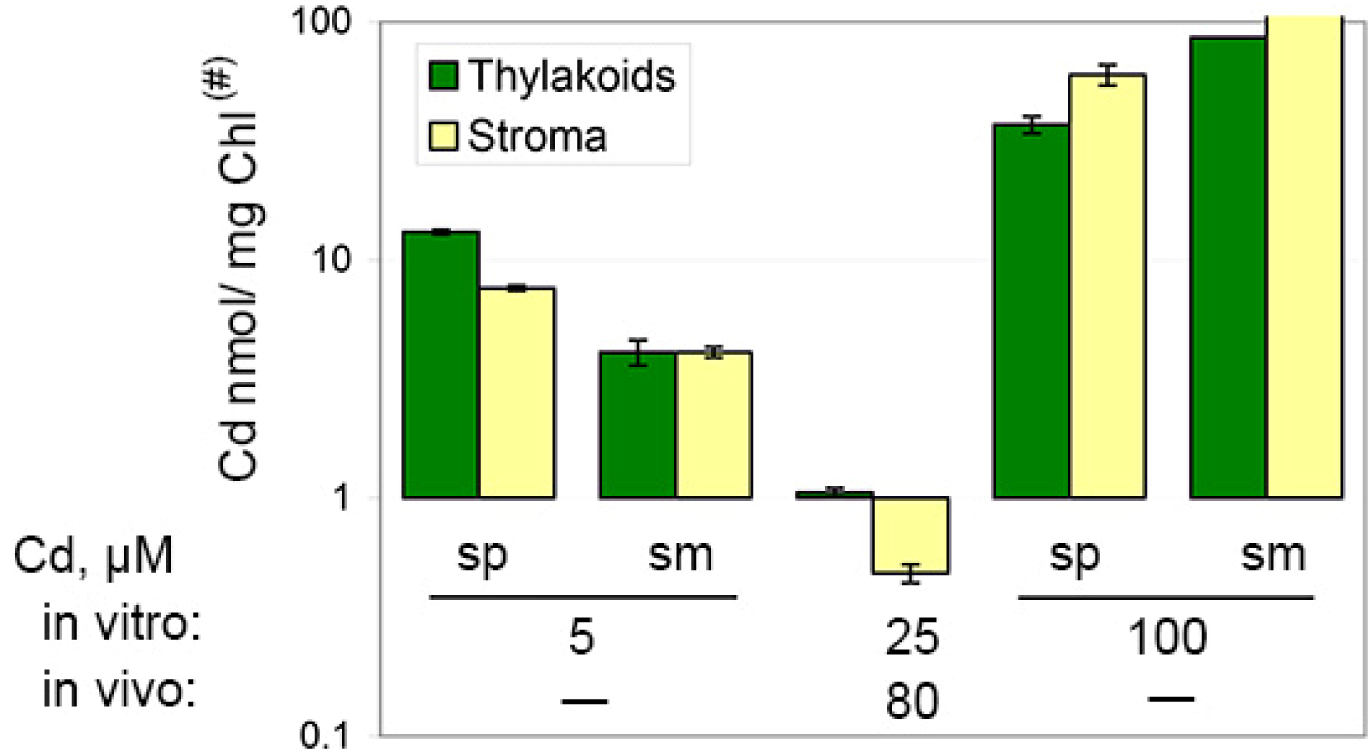
Cd accumulation *in vitro* by chloroplasts from untreated and Cd-treated barley plants Cd accumulation by chloroplasts from Cd-untreated plants is taken from Lysenko et al. (2019); Cd accumulation by chloroplast from Cd-treated plants is taken from Fig. 5. The both experiments were very similar (see text). Green – thylakoids, yellow – stroma+envelope. “sp” and “sm” – same experiments performed in spring and summer correspondingly. *In vivo*, plants were grown with no Cd addition (“-“) or at 80 μM Cd (“80”). *In vitro*, chloroplasts were incubated at 5, 25, or 100 μM Cd. # - designated as in Figs 3 and 5. Data are given as means ± SE.

Both experiments were nearly identical; chloroplasts were isolated from nine-day-old barley plants grown in the same conditions except Cd treatment; Cd accumulations *in vitro* were performed in Tris-based buffer for 75 min (Lysenko et al. 2019) or in Tricine-based buffer for 90 min (Fig. 5-6, Suppl. Table S9). Untreated barley plants demonstrated absence of Cd in the first and second leaves of 9- and 16-day-old plants (Lysenko et al. 2015, 2026), while low Cd content determined in roots and stems of these plants (Lysenko et al. 2015) and in leaves and other organs of similarly grown untreated maize plants (Klaus et al. 2013; Lysenko et al. 2015). Apparently, Cd accumulation in leaves of Cd-treated barley plants induced an unknown protecting mechanism that greatly reduced Cd transport across chloroplast envelope membranes. Such protective mechanism was not discussed previously.

### 4.3 Cd loss from chloroplasts

*In vivo*, thylakoids lost 17% or 34% of initial Cd and acquired 30% or 24% of late Cd (“uncorrected % or corrected %” values are given; Fig. 3A-C). *In vitro*, thylakoids lost 50% of initial Cd and accumulated portion of late Cd that was similar (108%) to the initial Cd content (Fig. 5A-B). *In vivo*, stroma and envelope lost 22% or 39% of initial Cd and acquired 22% or 17% of late Cd (“uncorrected % or corrected %” values are given; Fig. 3D-F). *In vitro*, stroma and envelope showed no loss of initial Cd and accumulated four times larger portion of late Cd (Fig. 5C-D). Therefore, we dealt with two different mechanisms. A mechanism of Cd removal from thylakoids functioned in both intact plants and isolated chloroplasts. A mechanism of Cd removal from stroma and envelope worked in intact plants but did not work in isolated chloroplasts.

A mechanism of Cd removal from thylakoids appeared to function much more effectively in isolated chloroplasts compared to intact plants. But why? Probably, Cd removal from chloroplasts *in vivo* was masked by some circumstances. First, initial Cd continued translocation from roots to shoot after 8^th^ day when initial Cd was replaced by late Cd in the mineral medium. After 8^th^ day, the first leaf accumulated 6.0 ± 0.45 nmol of late Cd (Fig. 2L-M); the amount of initial Cd transported to all leaves can be estimated as 9 ± 4 nmol. Likely, initial Cd was simultaneously transported into and out of the first leaf and both portions were equal (see chapter 4.1). Barley and maize accumulated more Cd in lower leaves and less Cd in leaves of higher storeys (Klaus et al. 2013; Lysenko et al. 2015). Thus, we can suppose that equal portions (∼6 nmol) of initial Cd and late Cd were translocated to the first leaf from 8^th^ to 16^th^ day; amount of initial Cd could be even larger. Therefore, we can admit that equal portions of initial Cd and late Cd were introduced to chloroplasts after 8^th^ day. In chloroplasts of 16-day-old plants, we can distinguish 23% or 29% (0.3 or 0.4 nmol/mg Chl depending on the correction) of late Cd due to difference between isotopes; however, we cannot recognize an equal portion (“23-29%”) of initial Cd acquired *de novo* because initial Cd accumulated before and after 8^th^ day were represented with the same isotope. Next, we should conclude that chloroplasts lost additional “23-29%” of initial Cd which was substituted with new uptake of “23-29%” of initial Cd transported from roots. This portion of removed Cd cannot be differentiated by isotope content.

Second, Chl content in 16-day-old leaves decreased by 20-25% compared to Chl content in 8-day-old leaves (Suppl. Table S5); probably, this was caused by corresponding Chl decrease in chloroplasts. A decrease of Chl content by 20-25% increases Cd/Chl ratio by 25-33% with no change of Cd. In chloroplasts, Cd was calculated per mg Chl solely; therefore, longer speculations are needed to show this effect. For example, the first leaves of 8-16 days plants (Suppl. Table S3, Fig. 2) and of cut shoots (Suppl. Table S8) contained similar amount of Cd per DW or per organ. Of all Cd in the first leaves, chloroplasts accumulated 1-2% in 8-16 days plants (Fig. 4F, Lysenko et al. 2015, 2019) and 1.62% in cut shoots (Suppl. Table S8). Therefore, chloroplasts contained similar amounts of Cd in both intact plants and cut shoots. However, threefold difference was observed per mg Chl. Cd content in chloroplasts was 1.36-1.51 nmol/mg Chl in 8-16 days plants (Suppl. Table S7) and 4.13 nmol/mg Chl in cut shoots (Suppl. Table S8, Fig. 4). This great difference was achieved due to severe Chl degradation in leaves and chloroplasts of cut shoots.

We calculated correction(s) to eliminate the impact of Chl decrease on Cd content in chloroplasts of 16-day-old plants (Eq. 4, Eq. 8). Corrected values revealed larger range of Cd removal from chloroplasts (Fig. 3). This range should be even bigger assuming *de novo* uptake of initial Cd by chloroplasts discussed above. *In vivo*, removal of bulk of Cd from thylakoids could be performed as effectively as it was observed *in vitro* where this process was not masked.

### 4.4 Possible mechanisms

To hypothesize possible mechanism(s), we should understand three important points. Inside chloroplasts, the most of Cd is located in thylakoids. Plants have two distinct mechanisms for Cd removal from thylakoids and stroma. In both stroma and thylakoids, the most of Cd is tightly bound. The first two points were discussed above; the latter point resulted from our experiments with EDTA. When EDTA was included to the incubation buffer, isolated chloroplasts did not accumulated Cd *in vitro* (Lysenko et al. 2019); probably, Cd was firmly bound by EDTA. When EDTA was included to the post-washing buffer, it removed 80-90% of freshly accumulated Cd from both stoma and thylakoids of intact chloroplasts; however, essential cations Mg, Fe, and K were not removed by EDTA (Lysenko et al. 2019). We concluded that the bivalent cations accumulated *in vivo* (Mg and Fe) were tightly bound while Cd accumulated for 75 min *in vitro* was mostly unbound or loosely bound. Therefore, the bulk of freshly accumulated Cd was removed by EDTA. The present study supported this conclusion. Cd accumulated *in vivo* was resistant to EDTA post-washing (Fig. 5). An alternative recalculation revealed small significant EDTA-dependent decrease (∼5%) of Cd that was accumulated by thylakoids *in vivo* in experiment ^114^Cd/^111^Cd; in combined data of both reciprocal experiments, this decrease was insignificant (Fig. 5E). The general resolution of our experimental approach is about 15%. The revealed 5% decrease can be false significant. Otherwise, Cd accumulated by thylakoids *in vivo* can include small portion (≤ 5%) of unbound and/or loosely bound Cd and we were succeeded to reveal it by chance. The contents of essential metals accumulated *in vivo* were mainly unchanged by EDTA-post-washing. The levels of Mg and Cu in thylakoids and stroma, Fe in thylakoids, and K in stroma were not influenced by EDTA (Fig. 6, Suppl. Table S9); data for K in thylakoids and Fe in stroma were unreliable, while Mn changes will be discussed below. Chloroplasts isolated from Cd-treated plants accumulated *in vitro* drastically smaller portion of Cd than chloroplasts from untreated plants did (Fig. 8). Of this small amount of freshly accumulated Cd, EDTA post-washing removed 17.3 ± 3.5% and 12.6 ± 5.0% of Cd from thylakoids and stroma correspondingly (Fig. 5E-F). Therefore, 80-90% of small portion of Cd in chloroplasts were tightly bound even after 90 min accumulation *in vitro*.

Zhao with colleagues demonstrated that inactivation of *Sedum plumbizincicola* gene HMA1 increased Cd content in chloroplasts; therefore, they concluded that SpHMA1 exports Cd from chloroplasts in this Cd-hyperaccumulator plant species (Zhao et al. 2019). However, studies of HMA1 function originated controversial results. HMA gene family encodes P_1B_-type ATPases (Axelsen and Palmgren 2001; Williams and Mills 2005). Clear orthologs of HMA1 were found in different dicotyledonous and monocotyledonous species (Huang et al. 2022; Zang et al. 2023; Shao et al. 2025). HMA1 is located in chloroplasts; it was shown for *A. thaliana* (Seigneurin- Berny et al. 2006; Kim et al. 2009). *S. plumbizincicola* (Zhao et al. 2019), and in heterologous system when barley gene was expressed in tobacco leaves (Mikkelsen et al. 2012). However, when HMA1 of *Capsicum annuum* was expressed in protoplasts of *A. thaliana*, then it was observed in endoplasmic reticulum and Golgi apparatus (Xu et al. 2023). In arabidopsis, HMA1 is located in chloroplast envelope (Seigneurin-Berny et al. 2006; Kim et al. 2009). Heterologous expression in yeast demonstrated that HMA1 can participate in transport of many bivalent cations. A role in Cu, Zn, and Cd transport was demonstrated with the genes of arabidopsis (Seigneurin-Berny et al. 2006; Moreno et al. 2008; Kim et al. 2009), *S. plumbizincicola* (Zhao et al. 2019, 2025; Huang et al. 2022), and barley (Mikkelsen et al. 2012). Arabidopsis and barley genes confirmed HMA1 participation in Ca transport (Moreno et al. 2008; Mikkelsen et al. 2012). Barley HMA1 demonstrated a role in Co, Mn, and Fe transport (Mikkelsen et al. 2012), while HMA1 of *S. plumbizincicola* showed no role in transport of Co and Mn (Zhao et al. 2025). At 70-80 μM Cd, growth of Cd-sensitive yeast strain *Δycf1* was inhibited by CaHMA1 (Xu et al. 2023) and stimulated by AtHMA1 (Moreno et al. 2008). Instead, Cu solely increased general ATPase activity in envelope of arabidopsis chloroplasts whereas Zn, Co, Ag, Mn, and Fe did not influence it (Seigneurin-Berny et al. 2006). General ATPase activity decreased Cu and Zn contents in barley chloroplasts, while Fe content remained unchanged (Mikkelsen et al. 2012); in these barley chloroplasts, Fe content was similar to ours while even decreased Cu and Zn contents exceeded our results several times (Lysenko et al. 2019, Suppl. Tables 7 and 9). Under normal conditions, knockout of AtHMA1 decreased in chloroplasts Cu content and total SOD activity, while Zn content did not change (Seigneurin-Berny et al. 2006). Under excess of Zn, knockout of AtHMA1 increased in chloroplasts Zn content by 30-40%, while Cu content remained unchanged (Kim et al. 2009). Arabidopsis chloroplasts bear two more members of this family – AtHMA6 (Shikanai et al. 2003) and AtHMA8 (Abdel-Ghany et al. 2005). Knockout of AtHMA6 decreased Cu content in chloroplasts (Shikanai et al. 2003; Boutigny et al. 2014); in ΔAtHMA6 mutant, overexpression of AtHMA1 restored Cu content in chloroplasts to WT level (Boutigny et al. 2014). Solely, knockout of HMA1 in *S. plumbizincicola* increased Cd content in chloroplasts (Zhao et al. 2019). Thus, we are far from understanding HMA1 role in chloroplasts. Likely, HMA1 performs transport of Cu into chloroplasts and can translocate other bivalent cations (Zn, Cd and other) in same direction in excluder species such as arabidopsis and barley. In *S. plumbizincicola,* HMA1 could have minimal affinity to Cd and translocates into chloroplasts essential cations like Cu and Zn with smallest Cd addition; in the absence of HMA1, its function can be substituted with other transporters with higher affinity to Cd. In recent articles, researchers of *S. plumbizincicola* support the opinion that SpHMA1 remove Cd from chloroplasts (Huang et al. 2022), whereas researchers of excluder plant species have not gone beyond simple statement that SpHMA1 someway decreased Cd content in chloroplasts (Xu et al. 2023; Zang et al. 2023; Shao et al. 2025). In any case, envelope transporter cannot explain why bulk of Cd was removed from thylakoids whereas Cd from stroma and envelope was not excluded *in vitro* (Fig. 5). However, it can be explained by versatile system of vesicular traffic.

Plants own two vesicular mechanisms for degradation of damaged cell structures or under senescence. In microautophagy, a cell structure (“cargo”) is surrounded by invagination or protrusion of tonoplast. In macroautophagy, a cargo is surrounded by double-membrane and transported to vacuole; this double-membrane is called autophagosome and its formation depends on ATG (autophagy) genes (Liu and Basscham 2012). Inside vacuole, cargos are degraded. Both micro- and macroautophagy are implicated in degradation of chloroplasts that is called chlorophagy. Chlorophagy machinery reviewed in (Soto-Burgos et al. 2018; Zhuang and Jiang 2019). Chlorophagy depends on ATG genes 5-8 and generates vesicles containing components of stroma or thylakoids. Vesicles comprised stromal content are represented with Rubisco containing bodies (RCB, Chiba et al. 2003), small starch granule-like (SSGL, Wang et al. 2013) structures, and senescence associated vacuoles (SAVs, Martinez et al. 2008).

Thylakoid-containing vesicles are represented with ATI-PS-bodies (ATG8-interacting- photosystem-bodies, Michaeli et al. 2014) and CV-containing vesicles (CCV, Wang and Blumwald, 2014). Whole chloroplasts can be covered by autophagosome membrane and translocated to vacuole (Wada et al. 2009; Izumi et al. 2017). Proteins ATI 1 and 2 (Michaeli et al. 2014) and VIPP1 (vesicle inducing protein in plastids 1, Nakamura et al. 2018), are located on chloroplast envelope; protein CV (chloroplast vesiculation) is located inside chloroplasts and interacts with PsbO subunit of photosystem II (Wang and Blumwald, 2014); NBR (neighbor of BRCA1) 1 protein is located on the surface and inside chloroplasts (Lee et al. 2023). Cd treatment stimulate autophagy system in *A. thaliana* (Wu et al. 2025). Vesicular translocation of thylakoids to vacuole can be responsible for well-known effect of Chl decrease in leaves of Cd- treated plants (*e.g.*, Suppl. Table S5, Lysenko et al. 2015); Cd content in chloroplasts is too low for a direct impairment of Chl molecules (discussed in Lysenko et al. 2015).

Therefore, chlorophagy machinery can independently remove large pieces of thylakoids or stroma; protein complexes with tightly bound Cd can be removed by these distinct pathways. Some of these pathways could be performed in isolated chloroplasts, while others cannot (*e.g.*, requires vacuoles).

### 4.5 EDTA effect on Mn

EDTA is well-known chelator of bivalent cations (*e.g.*, Sawyer and Paulsen 1959). EDTA post-washing did not change contents of endogenously accumulated metals with a sole exception. In chloroplasts isolated from Cd-treated plants, EDTA post-washing released a portion (∼14%) of Mn from thylakoids to stroma whereas total chloroplast content did not change significantly (Fig. 6B). This pattern observed in both reciprocal variants; Mn decrease in thylakoids was significant in each reciprocal variant, while Mn increase in stroma was only significant in combined values of eight independent experiments (Suppl. Table S9). Similar tendency was observed in chloroplasts isolated from untreated plants (Lysenko et al. 2019); these data were less undoubtful and were omitted then while we have shown them in Suppl. Table S10. The previous experiment was performed in two repeats only and *in vitro* reaction carried out in Tris buffer. Tris has ability to chelate bivalent cations (Fisher et al. 1979); Tris in high concentration (0.8 M) is applied to remove Mn from oxygen-evolving complex (OEC) of photosystem II (*e.g.*, Klimov et al. 1982). Therefore, in the current *in vitro* experiment Tris was replaced by Tricine.

We supposed that 2 mM EDTA treatment released Mn from Mn_4_CaO_5_-cluster of OEC in intact chloroplasts isolated from both untreated (Suppl. Table S10) and Cd-treated plants (Fig. 6B, Suppl. Table S9). However, OEC is rather resistant to EDTA and should be destabilized prior to a chelator application. 50 mM EDTA treatment of purified photosystem II did not changed OEC substantially; solely PsbU subunit was removed (Zhang et al. 2017). Pretreatment with 20-50 mM NH_2_OH reduced Mn in Mn_4_CaO_5_ to Mn^2+^; next, 50 mM EDTA washed out Mn and Ca completely (Zhang et al. 2017). Simultaneous washing with NH_2_OH and EDTA removed the most (∼95%) of Mn from photosystem II enriched membranes (Chen et al. 1995). High salt treatment also improved effect of EDTA (Kimura and Ono, 2003) or Tris (Klimov et al. 1982) on photosystem II enriched membranes. We washed intact chloroplasts with EDTA only. Chloroplasts contain reductants in stroma that could substitute NH_2_OH role. Damaged photosystem II complexes migrate to unstacked regions at thylakoid margins for reparation (Yoshioka-Nishimura 2016). Possibly, Mn_4_CaO_5_-clusters in photosystems II under reparation were reduced with stromal reductants and disintegrated with EDTA completely. However, we did not observe EDTA-induced release of initial Cd from thylakoids to stroma (Fig. 5). Therefore, a portion of disintegrated OEC (∼14%) was Cd-free. It contradicts to the conclusion that Cd substitute Ca or Mn in OEC *in vivo* (Bazzaz and Govinjee 1974; Sigfridson et al. 2004). Alternatively, stromal reductants and further chelation removed a single (particular?) Mn from a half of OEC.

Interestingly, EDTA treatment released Mn accumulated *in vivo* (Fig. 6B) and Cd accumulated *in vitro* (Fig. 5, Suppl. Tables S9 and S10) from thylakoids. The EDTA-released Cd disappeared from chloroplasts, while EDTA-released Mn retained in stroma. Probably, EDTA- released Cd was transported through envelope whereas Mn was not.

## 5. Conclusion

We advanced a model with two isotopes ^111^Cd and ^114^Cd. Applying this model to barley plants, we demonstrated several features. After a primary absorption by roots, Cd continued translocation to shoot for many days; a root barrier slowed down Cd translocation but did not halt it. In the process of growth, chloroplasts acquired new portions of Cd and lost part of Cd accumulated earlier. This loss of Cd demonstrated that excluder plant species has ability to remove Cd from chloroplasts. Removal of Cd from thylakoids was observed both *in vivo* and *in vitro*. Removal of Cd from stroma and envelope was carried out *in vivo* only; isolated chloroplasts were unable to remove Cd from stroma and envelope *in vitro*. Therefore, barley has distinct mechanisms for Cd removal from chloroplasts: one from thylakoids and another from stroma and envelope. Cd removal from chloroplast can be performed with the system of vesicular transport that is called chlorophagy. Diverse pathways of chlorophagy selectively transported fragments of thylakoids or stroma to vacuoles. The process of Cd accumulation by chloroplasts was mainly light-independent. *In vivo*, Cd in chloroplasts was tightly bound and mostly located in thylakoids. *In vitro*, chloroplasts isolated from Cd-treated plants accumulated several times smaller amount of Cd compared to chloroplasts from untreated plants. Probably, Cd-treated plants greatly reprogrammed transport across chloroplast envelope.

## Supplementary Information

The online version contains supplementary material.

## Author contribution

**Eugene A. Lysenko**: Conceptualization, Methodology, Project administration, Investigation, Visualization, Formal Analysis, Funding Acquisition, Writing; **Irina F. Seregina**: Investigation (measurement of ^111^Cd and ^114^Cd), **Alexander A. Klaus:** Investigation, Formal Analysis; **Alexander V. Kartashov**: Investigation. All authors read and approved the manuscript. EAL is the author responsible for contact and ensures communication.

## Supporting information

Supplemental Tables 6_7 complete

## Acknowledgements

Authors are grateful to Prof. G.A. Tsirlina for bridge building between the biologists and chemists, Drs. A.G. Borzenko and A.V. Shitov for helpful discussion, and N.A. Savvina for help in manuscript preparation. The research supported by the Russian Science Foundation grant №25-24-00009.

## Compliance with Ethical Standards

The authors declare no conflict of interest.

The authors have no relevant financial or non-financial interests to disclose.

Ethics declaration (Human/animal participants and informed consent): not applicable.

## Funding

The research supported by the Russian Science Foundation grant №25-24-00009. The funding source had no influence on the research process and manuscript preparation.

## Abbreviations

Chl –: chlorophyll
DW–: dry weight
FW –: fresh weight
HMA –: heavy metal ATPase of P_1B_-type
OEC –: oxygen-evolving complex of photosystem II
SE –: standard error

## Statements and Declarations

### Compliance with Ethical Standards

The authors declare no conflict of interest.

The authors have no relevant financial or non-financial interests to disclose.

Ethics declaration (Human/animal participants and informed consent): not applicable.

### Funding

This research was supported by the Russian Science Foundation grant №25-24-00009.

The funding source had no influence on the research process and manuscript preparation.

### Data Availability statement

The datasets generated and analysed during the current study are available from the corresponding author on reasonable request.

## Notes

### Competing Interest Statement

The authors have declared no competing interest.

### Summary of Updates

We have performed a correction to Cd contents presented in Fig. 3. We have added corrected values to Fig. 3 and extended this figure. This makes Fig. 9 obsolete and we have removed it. We have added equations (2-8) used for the corrections to Methods. We have added a couple of new speculations in Discussion and rewritten it more clearly.

