## Supplemental Tables 1_10 for "Common excluder barley has more than one mechanism to remove Cd from chloroplasts"

**Source:** bioRxiv

Table S1. Conditions of ICP-MS determination.

| | Parameters | Conditions |
|---|---|---|
| Plasma: | RF power | 1500 W |
| Gas flow rate: | Plasma gas | $15\ L\cdot\ min^{-1}$ |
| | Auxiliary gas | $1.2\ L\cdot\ min^{-1}$ |
| Data acquisition: | Dwell time | 0.1–0.5 s |
| | Number of replicates | 3 |
| Mass-spectrometer: | Resolution | 0.7 amu |
| | Vacuum without plasma | $4\times10^{-5}$ Torr |
| | Vacuum with plasma | $4\times10^{-4}$ Torr |

Table S2. Changes of the first leaves of young barley plants during six days at Cd-free mineral medium.

| Parameters | | Cd treatment + post-treatment time (days) | | | |
|---|---|---|---|---|---|
| | | 9d | 9d + 2d | 9d + 4d | 9d + 6d |
| FW | mg | 89.6 ± 2.9 | 89.5 ± 2.5 [a] | 95.9 ± 3.1 | 96.5 ± 2.2 [b] |
| DW | mg | 8.55 ± 0.31 | 8.32 ± 0.15 | 8.40 ± 0.35 | 8.34 ± 0.19 |
| $H_2O$ content | % | 90.5 ± 0.2 [a] | 90.7 ± 0.1 [a] | 91.3 ± 0.1 [b] | 91.3 ± 0.2 [b] |
| Chl a | µg/g FW | 983 ± 71 | 940 ± 56 | 896 ± 104 | 827 ± 137 |
| Chl b | µg/g FW | 287 ± 27 | 276 ± 22 | 267 ± 36 | 250 ± 44 |
| Car | µg/g FW | 150 ± 8 | 145 ± 7 | 132 ± 11 | 127 ± 14 |
| Chl a/b | | 3.45 ± 0.03 [a] | 3.43 ± 0.03 [a] | 3.37 ± 0.03 | 3.27 ± 0.04 [b] |
| Chl/Car | | 8.64 ± 0.35 | 8.41 ± 0.35 | 8.25 ± 0.45 | 7.93 ± 0.48 |
| Chl/Car [§] | [9d = 1] [§] | 1 ± 0 [a] | 0.97 ± 0.01 | 0.95 ± 0.02 | 0.92 ± 0.03 [b] |

Until ninth day, plants were grown on Cd containing medium (80 µM); next, plants were transferred to fresh portion of the medium with no Cd addition and grown for another six days. Data are given as means ± SE. a-b – significantly different values, $p < 0.05$; no lettering means absence of significant differences (to ease perception). First leaf – leaf blade.
[§] – in each repeat of the experiment, variant 9d accepted as 1 and other three variants calculated as portion of 1.

*Probably, the increase of water content is responsible for the increase of FW and the decrease of Chls and carotenoids per g FW.*

Table S3. Changes of metal contents in the first leaves and chloroplasts during six days at Cd-free mineral medium.

| Me / variant | Metal contents | | | |
|---|---|---|---|---|
| | First leaves | | Chloroplasts in first leaves | |
| | μmol/g DW | nmol per a leaf | nmol/mg Chl | % from leaves [×] |
| Cd | | | | |
| 9d | 1.70 ± 0.15 | 14.5 ± 1.4 | 1.79 ± 0.19 | 1.40 ± 0.11 |
| 9d + 2d | 1.82 ± 0.14 | 15.1 ± 1.0 | 1.96 ± 0.16 | 1.39 ± 0.13 |
| 9d + 4d | 2.08 ± 0.33 | 17.1 ± 2.3 | 2.17 ± 0.48 | 1.21 ± 0.18 |
| 9d + 6d | 2.13 ± 0.30 | 17.5 ± 2.0 | 2.54 ± 0.54 | 1.38 ± 0.34 |
| Cu | | | | |
| 9d | 0.050 ± 0.004 [a] | 0.434 ± 0.050 | 1.56 ± 0.09 [a] | 42 ± 5 |
| 9d + 2d | 0.044 ± 0.007 | 0.368 ± 0.068 | 1.51 ± 0.20 | 46 ± 7 |
| 9d + 4d | 0.040 ± 0.004 | 0.340 ± 0.044 | 1.27 ± 0.15 | 37 ± 4 |
| 9d + 6d | 0.035 ± 0.005 [b] | 0.293 ± 0.048 | 1.19 ± 0.07 [b] | 45 ± 4 |
| Mn | | | | |
| 9d | 0.632 ± 0.019 | 5.41 ± 0.31 | 8.50 ± 0.15 | 18 ± 1 |
| 9d + 2d | 0.662 ± 0.028 | 5.52 ± 0.33 | 8.28 ± 0.41 | 17 ± 3 |
| 9d + 4d | 0.687 ± 0.029 | 5.78 ± 0.42 | 7.80 ± 0.85 | 13 ± 2 |
| 9d + 6d | 0.685 ± 0.017 | 5.70 ± 0.13 | 8.05 ± 0.29 | 17 ± 3 |
| Fe | | | | |
| 9d | 1.56 ± 0.09 | 13.3 ± 0.7 | 66.3 ± 5.7 | 58 ± 8 |
| 9d + 2d | 1.68 ± 0.11 | 13.7 ± 0.9 | 67.6 ± 9.6 | 49 ± 16 |
| 9d + 4d | 1.48 ± 0.15 | 12.5 ± 1.6 | 59.0 ± 5.7 | 47 ± 5 |
| 9d + 6d | 1.37 ± 0.13 | 11.5 ± 1.4 | 68.3 ± 2.8 | 63 ± 16 |
| K | | | | |
| 9d | 1477 ± 74 | 12 558 ± 592 [a] | 17.1 ± 0.8 [a] | 0.016 ± 0.002 [a] |
| 9d + 2d | 1486 ± 46 [a] | 12 355 ± 395 [a] | 24.8 ± 2.9 [a] | 0.023 ± 0.004 |
| 9d + 4d | 1706 ± 81 [b] | 14 224 ± 385 [b] | 48.7 ± 14.1 | 0.033 ± 0.010 |
| 9d + 6d | 1560 ± 74 | 12 994 ± 667 | 40.5 ± 5.1 [b] | 0.030 ± 0.005 [b] |
| Zn | | | | |
| 9d | 0.565 ± 0.119 | 4.87 ± 1.11 | - | - |
| 9d + 2d | 0.728 ± 0.226 | 6.04 ± 1.89 | - | - |
| 9d + 4d | 0.905 ± 0.535 | 7.52 ± 4.30 | - | - |
| 9d + 6d | 0.748 ± 0.490 | 6.07 ± 3.87 | - | - |
| Ca | | | | |
| 9d | 77 ± 2 [a] | 658 ± 30 [a] | - | - |
| 9d + 2d | 102 ± 7 [b] | 850 ± 65 [b] | - | - |
| 9d + 4d | 125 ± 9 [b] | 1047 ± 69 [b] | - | - |
| 9d + 6d | 121 ± 17 [b] | 1000 ± 122 [b] | - | - |
| Na | | | | |
| 9d | 115 ± 16 [a] | 991 ± 158 [a] | - | - |
| 9d + 2d | 173 ± 25 | 1449 ± 237 | - | - |
| 9d + 4d | 259 ± 30 [b] | 2155 ± 212 [b] | - | - |
| 9d + 6d | 218 ± 42 [b] | 1796 ± 313 [b] | - | - |

Designations are the same as in Table S2.

[×] – portion of Cd in chloroplasts (%) of all Cd in the first leaves calculated by equation: $C/L \times 100\%$, where C is the Cd content in chloroplasts (nmol/mg Chl), and L is the Cd content in leaves (nmol/mg Chl).

Table S4. Changes of organs of young barley plants during eight days at Cd-containing mineral medium.

| Organ | Day | FW, mg | | DW, mg | | $H_2O$ content, % | |
|---|---|---|---|---|---|---|---|
| | | $^{111}$Cd/ $^{114}$Cd | $^{114}$Cd/ $^{111}$Cd | $^{111}$Cd/ $^{114}$Cd | $^{114}$Cd/ $^{111}$Cd | $^{111}$Cd/ $^{114}$Cd | $^{114}$Cd/ $^{111}$Cd |
| First leaf | 8 | 103 ± 2 $^{a}$ | 101 ± 3 $^{a}$ | 11.0 ± 0.3 $^{a}$ | 10.9 ± 0.2 $^{a}$ | 89.3 ± 0.3 $^{a}$ | 89.2 ± 0.2 $^{a}$ |
| | 12 | 101 ± 3 $^{a}$ | 107 ± 4 $^{a}$ | 11.8 ± 0.5 $^{a}$ | 12.3 ± 0.2 $^{b}$ | 88.3 ± 0.2 $^{b}$ | 88.5 ±0.2 $^{b}$ |
| | 16 | 88 ± 2 $^{b}$ * | 98 ± 4 $^{a}$ * | 10.7 ± 0.3 $^{a}$ | 11.3 ± 0.5$^{ab}$ | 87.8 ± 0.2 $^{b}$ | 88.5 ±0.4$^{ab}$ |
| Shoot except first leaf | 8 | 59 ± 2 $^{a}$ | 55 ± 2 $^{a}$ | 6.5 ± 0.5 $^{a}$ | 6.1 ± 0.1 $^{a}$ | 88.9 ± 0.4 $^{a}$ | 88.8 ± 0.2 $^{a}$ |
| | 12 | 139 ± 5 $^{b}$ | 149 ± 6 $^{b}$ | 16.3 ± 0.8 $^{b}$ | 17.3 ± 1.4 $^{b}$ | 88.3 ± 0.3 $^{a}$ | 88.4 ± 0.2$^{ab}$ |
| | 16 | 198 ± 7 $^{c}$ | 207 ± 8 $^{c}$ | 26.2 ± 1.4 $^{c}$ | 26.3 ± 1.7 $^{c}$ | 86.9 ± 0.3 $^{b}$ | 87.4 ± 0.4 $^{b}$ |
| Roots | 8 | 43 ± 2 $^{a}$ | 43 ± 2 $^{a}$ | 5.25 ± 0.20 $^{a}$ | 5.38 ± 0.16 $^{a}$ | 87.8 ± 0.6 $^{a}$ | 87.5 ± 0.3 $^{a}$ |
| | 12 | 58 ± 3 $^{b}$ | 55 ± 4 $^{b}$ | 7.24 ± 0.37 $^{b}$ | 7.41 ± 0.42 $^{b}$ | 87.5 ± 0.3 $^{a}$ | 87.7 ± 0.6 $^{a}$ |
| | 16 | 61 ± 5 $^{b}$ | 63 ± 6 $^{b}$ | 8.87 ± 0.51 $^{c}$ | 8.26 ± 0.66 $^{b}$ | 87.2 ± 0.5 $^{a}$ | 86.8 ± 0.8 $^{a}$ |

Barley plants were grown in the presence of one isotope ($^{111}$Cd or $^{114}$Cd) from 3$^{rd}$ to 8$^{th}$ day; next, they were transferred to the alternative Cd isotope and grown until 16$^{th}$ day. The treatments with "a first" (initial) and "a second" (late) Cd performed in two reciprocal variants: $^{111}$Cd/ $^{114}$Cd and $^{114}$Cd/ $^{111}$Cd.

First leaf – leaf blade; roots – of a single plant.

a-c – significant differences between plants of diverse ages in a single variant, $p < 0.05$.

* – significant difference between reciprocal variants ($^{111}$Cd/ $^{114}$Cd and $^{114}$Cd/ $^{111}$Cd), $p < 0.05$.

Table S5. Changes of Chl and carotenoids (Car) in the first leaves of barley plants during eight days at Cd-containing mineral medium.

| Variant | Day | First leaves | | | | | Chloroplasts in first leaves | |
|---|---|---|---|---|---|---|---|---|
| | | μg/ g FW | | | | | | |
| | | Chl a | Chl b | Car | Chl a/b | Chl/Car | Chl a/b | Chl/Car |
| $^{111}$Cd/ $^{114}$Cd | 8 | 1184 ± 36 $^{a}$* | 359 ± 12 $^{a}$ | 195 ± 6 $^{a}$* | 3.30 ± 0.05 | 7.91 ± 0.17 | 3.30 ± 0.03 | 8.50 ± 0.19 |
| | 16 | 880 ± 41 $^{b}$ | 269 ± 14 $^{b}$ | 144 ± 8 $^{b}$ | 3.28 ± 0.08 | 8.02 ± 0.27 | 3.32 ± 0.06 | 8.45 ± 0.22 |
| $^{114}$Cd/ $^{111}$Cd | 8 | 1066 ± 26 $^{a}$* | 322 ± 14 $^{a}$ | 171 ± 2 $^{a}$* | 3.32 ± 0.08 | 8.10 ± 0.15 | 3.30 ± 0.01 | 8.49 ± 0.13 |
| | 16 | 877 ± 45 $^{b}$ | 269 ± 16 $^{b}$ | 146 ± 5 $^{b}$ | 3.27 ± 0.05 | 7.84 ± 0.25 | 3.31 ± 0.05 | 8.23 ± 0.15 |

Designations are the same as in Tables S2 and S4. Ratios Chl a/b and Chl/Car showed no significant difference and remained unmarked to ease perception.

Table S6 (reduced). Changes of metal contents in organs of barley plants during eight days at Cd-containing mineral medium.

| Me | Day | Organ | | | Me | Day | Organ | | |
|---|---|---|---|---|---|---|---|---|---|
| | | Roots | Stem+ | First leaf | | | Roots | Stem+ | First leaf |
| Cu | | nmol/ g DW | | | Mn | | nmol/ g DW | | |
| | 8 | 104 ± 8$^{ab}$ | 66.7 ± 5.0$^{a}$ | 33.2 ± 4.3$^{a}$ | | 8 | 373 ± 16$^{a}$ | 450 ± 19$^{a}$ | 554 ± 10$^{a}$ |
| | 12 | 105 ± 4$^{a}$ | 33.8 ± 3.8$^{b}$ | 25.4 ± 2.9$^{a}$ | | 12 | 287 ± 11$^{b}$ | 294 ± 8$^{b\ (c)}$ | 605 ± 16$^{b}$ |
| | 16 | 90 ± 3$^{b}$ | 27.8 ± 2.4$^{b}$ | 22.4 ± 2.3$^{a}$ | | 16 | 302 ± 9$^{b}$ | 309 ± 8$^{b}$ | 706 ± 12$^{c}$ |
| | | pmol/ organ | | | | | nmol/ organ | | |
| | 8 | 551 ± 45$^{a}$ | 421 ± 33$^{a}$ | 360 ± 45$^{a}$ | | 8 | 1.99 ± 0.12$^{a}$ | 2.83 ± 0.12$^{a}$ | 6.07 ± 0.15$^{a}$ |
| | 12 | 762 ± 28$^{b}$ | 572 ± 74$^{ab}$ | 304 ± 35$^{ab}$ | | 12 | 2.11 ± 0.13$^{a}$ | 4.89 ± 0.17$^{b}$ | 7.29 ± 0.24$^{b}$ |
| | 16 | 762 ± 28$^{b}$ | 725 ± 64$^{b}$ | 245 ± 24$^{b}$ | | 16 | 2.60 ± 0.16$^{b}$ | 8.11 ± 0.37$^{c}$ | 7.77 ± 0.20$^{b}$ |
| Zn | | µmol/ g DW | | | Fe | | µmol/ g DW | | |
| | 8 | 13.6 ± 2.5$^{a}$ | 1.90 ± 0.34$^{a}$ | 1.35 ± 0.30$^{a}$ | | 8 | 34.1 ± 3.9$^{a}$ | 1.07 ± 0.05$^{a}$ | 1.51 ± 0.07$^{a}$ |
| | 12 | 18.5 ± 1.2$^{a\ (b)}$ | 2.91 ± 0.13$^{b}$ | 1.47 ± 0.13$^{a}$ | | 12 | 82.1 ± 8.2$^{b}$ | 1.46 ± 0.11$^{b}$ | 1.95 ± 0.18$^{b}$ |
| | 16 | 26.5 ± 1.6$^{c}$ | 4.86 ± 0.46$^{c}$ | 2.03 ± 0.19$^{b}$ | | 16 | 108.1 ± 9.2$^{c}$ | 1.46 ± 0.02$^{b}$ | 2.01 ± 0.16$^{b}$ |
| | | nmol/ organ | | | | | nmol/ organ | | |
| | 8 | 72 ± 13$^{a}$ | 11.9 ± 2.1$^{a}$ | 14.6 ± 3.1$^{a}$ | | 8 | 182 ± 21$^{a}$ | 6.7 ± 0.3$^{a}$ | 16.6 ± 0.8$^{a}$ |
| | 12 | 135 ± 9$^{b}$ | 49.2 ± 4.0$^{b}$ | 17.6 ± 1.5$^{a}$ | | 12 | 599 ± 57$^{b}$ | 24.1 ± 1.5$^{b}$ | 23.7 ± 2.6$^{b}$ |
| | 16 | 230 ± 22$^{c}$ | 128.5 ± 13.6$^{c}$ | 22.5 ± 2.4$^{a}$ | | 16 | 949 ± 112$^{c}$ | 38.3 ± 1.8$^{c}$ | 22.4 ± 2.3$^{b}$ |
| Mg | | µmol/ g DW | | | Ca | | µmol/ g DW | | |
| | 8 | 61.5 ± 2.0$^{a}$ | 77.6 ± 1.5$^{a}$ | 64.5 ± 1.2$^{a}$ | | 8 | 63.6 ± 1.6$^{a}$ | 79 ± 3$^{a}$ | 132 ± 6$^{a}$ |
| | 12 | 53.8 ± 1.4$^{b}$ | 69.2 ± 2.8$^{b}$ | 59.0 ± 2.0$^{c}$ | | 12 | 76.8 ± 4.4$^{c\ (b)}$ | 76 ± 5$^{a}$ | 176 ± 7$^{b}$ |
| | 16 | 52.2 ± 1.6$^{b}$ | 72.5 ± 1.6$^{b}$ | 67.5 ± 1.8$^{a}$ | | 16 | 84.6 ± 2.1$^{c}$ | 107 ± 8$^{b}$ | 231 ± 8$^{c}$ |
| | | nmol/ organ | | | | | nmol/ organ | | |
| | 8 | 327 ± 13$^{a}$ | 489 ± 19$^{a}$ | 705 ± 14$^{a}$ | | 8 | 338 ± 11$^{a}$ | 501 ± 31$^{a}$ | 1439 ± 68$^{a}$ |
| | 12 | 394 ± 19$^{b}$ | 1144 ± 37$^{b}$ | 712 ± 30$^{a}$ | | 12 | 570 ± 52$^{b}$ | 1256 ± 80$^{b}$ | 2121 ± 96$^{b}$ |
| | 16 | 445 ± 21$^{b}$ | 1904 ± 92$^{c}$ | 741 ± 16$^{a}$ | | 16 | 731 ± 50$^{c}$ | 2847 ± 272$^{c}$ | 2540 ± 95$^{c}$ |
| Na | | µmol/ g DW | | | K | | µmol/ g DW | | |
| | 8 | 44.5 ± 2.7$^{a}$ | 104 ± 9$^{ab}$ | 129 ± 10$^{a}$ | | 8 | 774 ± 35$^{a}$ | 1278 ± 34$^{a}$ | 1655 ± 73$^{a}$ |
| | 12 | 34.2 ± 1.7$^{b}$ | 88 ± 6$^{a}$ | 148 ± 6$^{a}$ | | 12 | 797 ± 42$^{a}$ | 1704 ± 54$^{c\ (b)}$ | 1886 ± 58$^{c}$ |
| | 16 | 35.1 ± 2.0$^{b}$ | 110 ± 6$^{b}$ | 198 ± 10$^{c}$ | | 16 | 841 ± 19$^{a\ (b)}$ | 1844 ± 76$^{c}$ | 1641 ± 123$^{ac}$ |
| | | µmol/ organ | | | | | µmol/ organ | | |
| | 8 | 1.86 ± 0.07$^{a}$ | 0.66 ± 0.07$^{a}$ | 1.41 ± 0.11$^{a}$ | | 8 | 4.10 ± 0.19$^{a}$ | 8.04 ± 0.27$^{a}$ | 18.05 ± 0.71$^{a}$ |
| | 12 | 2.01 ± 0.10$^{ab}$ | 1.44 ± 0.06$^{b}$ | 1.78 ± 0.06$^{b}$ | | 12 | 5.86 ± 0.39$^{b}$ | 28.59 ± 1.63$^{b}$ | 22.82 ± 1.08$^{b}$ |
| | 16 | 2.30 ± 0.12$^{b}$ | 2.87 ± 0.19$^{c}$ | 2.18 ± 0.11$^{c}$ | | 16 | 7.24 ± 0.44$^{c}$ | 48.60 ± 3.15$^{c}$ | 18.17 ± 1.51$^{a}$ |

Complete version of the table is too large and represented in a separate file. The reduced version shows combined data of reciprocal pairs of experiments and omitted data per FW. No significant differences observed between reciprocal variants.
Me – metal; stem+ – stem with leaf sheaths. SI prefixes colored to ease perception.
Other designations are the same as in Tables S2 and S4. In parentheses – differences significant per FW.

Table S7 (reduced). Changes of metal contents in chloroplasts of barley first leaves during eight days at Cd-containing mineral medium.

| Metal | Day | Fractions of chloroplasts | | | | Chloroplasts | |
|---|---|---|---|---|---|---|---|
| | | Thylakoids | | Stroma + envelope | | nmol/mg Chl | % from leaves $^{\times}$ |
| | | nmol/mg Chl | % $^{\&}$ | nmol/mg Chl $^{\#}$ | % $^{\&}$ | | |
| Cd | 8 | 1.13 ± 0.02 $^{a}$ | 83.0 ± 0.8 | 0.23 ± 0.01 $^{a}$ | 17.0 ± 0.8 | 1.36 ± 0.03 $^{a}$ | 2.08 ± 0.07 $^{a}$ |
| | 16 | 1.28 ± 0.03 $^{b}$ | 84.6 ± 0.6 | 0.23 ± 0.01 $^{a}$ | 15.4 ± 0.6 | 1.51 ± 0.05 $^{b}$ | 1.09 ± 0.04 $^{b}$ |
| | 16^ | 1.01 ± 0.04 $^{c}$ | | 0.18 ± 0.01 $^{b}$ | | 1.20 ± 0.05 $^{c}$ | |
| Cu | 8 | 1.52 ± 0.07 $^{a}$ | 80.2 ± 1.8 | 0.38 ± 0.05 | 19.8 ± 1.8 | 1.90 ± 0.09 $^{a}$ | 94 ± 15 |
| | 16 | 1.07 ± 0.05 $^{b}$ | 74.8 ± 2.6 | 0.36 ± 0.04 | 25.2 ± 2.6 | 1.44 ± 0.05 $^{b}$ | 68 ± 7 |
| Mn | 8 | 9.8 ± 0.5 $^{a}$ | 86.4 ± 0.6 $^{a}$ | 1.51 ± 0.03 | 13.6 ± 0.6 $^{a}$ | 11.3 ± 0.5 $^{a}$ | 28.1 ± 1.8 $^{a}$ |
| | 16 | 8.3 ± 0.2 $^{b}$ | 84.5 ± 0.6 $^{b}$ | 1.52 ± 0.04 | 15.5 ± 0.6 $^{b}$ | 9.9 ± 0.2 $^{b}$ | 13.7 ± 0.5 $^{b}$ |
| Fe | 8 | 68.2 ± 2.7 | 95.4 ± 1.1 | 3.5 ± 1.0 | 4.6 ± 1.1 | 72.8 ± 3.1 | 67 ± 7 $^{a}$ |
| | 16 | 64.8 ± 4.1 | 95.1 ± 0.4 | 3.3 ± 0.3 | 4.9 ± 0.4 | 68.1 ± 4.3 | 35 ± 3 $^{b}$ |
| Mg | 8 | 1350 ± 73 | 98.5 ± 0.1 | 20.3 ± 1.6 | 1.5 ± 0.1 | 1370 ± 73 | 29.3 ± 2.3 $^{a}$ |
| | 16 | 1281 ± 27 | 98.7 ± 0.2 | 17.3 ± 2.7 | 1.3 ± 0.2 | 1298 ± 29 | 19.1 ± 1.0 $^{b}$ |
| | | | | | | | $ |
| K | 8 | n.d. | | 11.4 ± 1.3 $^{a}$ | | | 0.010 ± 0.002 |
| | 16 | n.d. | | 50.5 ± 13.8 $^{b}$ | | | 0.037 ± 0.013 |

Complete version of the table is large and represented in a separate file. The reduced version solely shows combined data of reciprocal pairs of experiments (initial Cd/ late Cd). No significant differences observed between reciprocal pairs of each variant.

The experiment is described in Table S4; chloroplasts isolated from the first leaves. Cd is shown as the sum of isotopes ($^{111}$Cd + $^{114}$Cd). $^{\&}$ – % in chloroplasts; $^{\#}$ – per mg Chl in chloroplasts from which the stromal fraction was separated; $^{\times}$ – see Table S3; $ – portion of K in stroma+envelope (%) of all K in the first leaves; n.d. – not detected. K content in thylakoids is tiny and required more chloroplasts for proper measurement (Lysenko et al. 2019). ^ - correction for the effect of Chl decrease; for the sum of isotopes, both corrections (Eq. 4 and Eq. 8) resulted in identical values and did not change proportions (%). a-b-c – significant difference between chloroplasts from 8- and 16-day-old plants in a corresponding variant, $p < 0.05$; pairs with no significant difference remained unmarked to ease perception. Differences between reciprocal variants were insignificant. Means ± SE.

Table S8. Metal contents in barley leaves and chloroplasts after reciprocal experiments *in organo*.

| Me | Variant | First leaves | | Chloroplasts | |
|---|---|---|---|---|---|
| | | nmol/ g DW | nmol/ organ | nmol/ mg Chl | % from leaves × |
| Cd | 111/ 114 | 1449 ± 145 | 16.8 ± 1.4 | 4.60 ± 0.52 | 1.66 ± 0.14 |
| | 114/ 111 | 1382 ± 106 | 16.3 ± 1.4 | 3.65 ± 0.39 | 1.57 ± 0.11 |
| | Σ | 1415 ± 84 | 16.6 ± 0.9 | 4.13 ± 0.35 | 1.62 ± 0.08 |
| Cu | 111/ 114 | 25.4 ± 1.6 | 0.300 ± 0.033 | 1.28 ± 0.09 | 27.7 ± 5.3 |
| | 114/ 111 | 26.4 ± 4.9 | 0.312 ± 0.059 | 1.22 ± 0.06 | 29.9 ± 4.5 |
| Mn | 111/ 114 | 651 ± 35 | 7.6 ± 0.4 | 8.3 ± 0.5 | 6.7 ± 0.8 |
| | 114/ 111 | 676 ± 35 | 8.0 ± 0.2 | 8.4 ± 0.3 | 7.5 ± 0.8 |
| Fe | 111/ 114 | 1241 ± 151 | 14.6 ± 2.0 | 195 ± 41 | 81 ± 12 |
| | 114/ 111 | 960 ± 45 | 11.3 ± 0.3 | 151 ± 26 | 99 ± 27 |
| Zn | 111/ 114 | 1630 ± 443 | 19.2 ± 5.2 | | |
| | 114/ 111 | 1829 ± 231 | 21.5 ± 2.6 | | |
| | | µmol/ g DW | µmol/ organ | | |
| Mg | 111/ 114 | 77 ± 5 | 0.90 ± 0.04 | 1216 ± 46 | 8.5 ± 1.1 |
| | 114/ 111 | 74 ± 2 | 0.87 ± 0.02 | 1299 ± 134 | 10.8 ± 2.1 |
| K | 111/ 114 | 1439 ± 192 | 16.6 ± 1.7 | 35.3 ± 17.1 | 0.014 ± 0.008 |
| | 114/ 111 | 1404 ± 81 | 16.5 ± 0.9 | 12.9 ± 2.1 | 0.006 ± 0.001 |
| Ca | 111/ 114 | 248 ± 24 | 2.9 ± 0.1 | | |
| | 114/ 111 | 235 ± 10 | 2.8 ± 0.2 | | |
| Na | 111/ 114 | 176 ± 31 | 2.0 ± 0.2 | | |
| | 114/ 111 | 196 ± 32 | 2.3 ± 0.3 | | |

Barley plants were grown at initial Cd ($^{111}$Cd or $^{114}$Cd, 80 µM) for five days (days 3-8); next, the shoots were cut and placed vertically on late Cd isotope ($^{114}$Cd or $^{111}$Cd, 30 µM) for six days (days 8-14). Chloroplasts isolated from the first (major) and second (minor) leaves. Cd is shown as the sum of isotopes ($^{111}$Cd + $^{114}$Cd). Σ – combined data of both reciprocal experiments. × – see Table S3. SI prefixes colored to ease perception. Differences between reciprocal variants were insignificant. Means ± SE.

Table S9. Changes of metal contents in barley chloroplasts during Cd accumulation *in vitro*.

| Variants | | Metal contents, nmol/mg Chl $^{(\#)}$ | | | Metal contents, nmol/mg Chl $^{(\#)}$ | | |
|---|---|---|---|---|---|---|---|
| | | Fractions of chloroplasts | | Chloroplasts | Fractions of chloroplasts | | Chloroplasts |
| *In vivo* | *In vitro* | Thylakoids | Stroma + | | Thylakoids | Stroma + | |
| | | Cd | | | Cu | | |
| $^{111}$Cd | - | 0.98 ± 0.08 $^{a}$ | 0.13 ± 0.02 $^{a}$ | 1.11 ± 0.09 $^{a}$ | 1.60 ± 0.28 $^{a}$ | 0.57 ± 0.05 $^{a}$ | 2.17 ± 0.28 $^{a}$ |
| | $^{114}$Cd | 1.52 ± 0.11 $^{b}$ | 0.59 ± 0.06 $^{b}$ | 2.11 ± 0.17 $^{b}$ | 1.40 ± 0.13 $^{a}$ | 0.46 ± 0.08 $^{a}$ | 1.87 ± 0.21 $^{a}$ |
| | $^{114}$Cd _E | 1.33 ± 0.06 $^{b}$ | 0.53 ± 0.05 $^{b}$ | 1.86 ± 0.05 $^{b}$ | 1.45 ± 0.23 $^{a}$ | 0.56 ± 0.13 $^{a}$ | 2.01 ± 0.34 $^{a}$ |
| $^{114}$Cd | - | 0.96 ± 0.05 $^{a}$ | 0.15 ± 0.02 $^{a}$ | 1.11 ± 0.07 $^{a}$ | 1.63 ± 0.34 $^{a}$ | 0.54 ± 0.09 $^{a}$ | 2.17 ± 0.43 $^{a}$ |
| | $^{111}$Cd | 1.53 ± 0.10 $^{b}$ | 0.62 ± 0.11 $^{b}$ | 2.16 ± 0.21 $^{b}$ | 1.42 ± 0.11 $^{a}$ | 0.47 ± 0.14 $^{a}$ | 1.89 ± 0.25 $^{a}$ |
| | $^{111}$Cd _E | 1.31 ± 0.07 $^{b}$ | 0.56 ± 0.09 $^{b}$ | 1.87 ± 0.16 $^{b}$ | 1.39 ± 0.08 $^{a}$ | 0.54 ± 0.15 $^{a}$ | 1.93 ± 0.18 $^{a}$ |
| 1$^{st}$ Cd | - | 0.97 ± 0.04 $^{a}$ | 0.14 ± 0.01 $^{a}$ | 1.11 ± 0.05 $^{a}$ | 1.61 ± 0.20 $^{a}$ | 0.56 ± 0.05 $^{a}$ | 2.17 ± 0.24 $^{a}$ |
| (Σ) | 2$^{nd}$ Cd | 1.53 ± 0.07 $^{b}$ | 0.61 ± 0.06 $^{b}$ | 2.13 ± 0.13 $^{b}$ | 1.41 ± 0.08 $^{a}$ | 0.47 ± 0.07 $^{a}$ | 1.88 ± 0.15 $^{a}$ |
| | 2$^{nd}$ Cd_E | 1.32 ± 0.04 $^{c}$ | 0.55 ± 0.05 $^{b}$ | 1.86 ± 0.08 $^{b}$ | 1.42 ± 0.11 $^{a}$ | 0.55 ± 0.09 $^{a}$ | 1.97 ± 0.18 $^{a}$ |
| | | Mn | | | Fe | | |
| $^{111}$Cd | - | 5.95 ± 0.38 $^{ab1}$ | 1.72 ± 0.29 $^{a}$ | 7.51 ± 0.78 $^{a}$ | 58.6 ± 5.6 $^{a}$ | 19.6 ± 9.5 $^{a}$ | 78.3 ±14.6 $^{a}$ |
| | $^{114}$Cd | 5.87 ± 0.12 $^{a}$ | 1.57 ± 0.16 $^{a}$ | 7.44 ± 0.19 $^{a}$ | 53.2 ± 2.7 $^{a}$ | 4.9 ± 2.0 $^{a}$ | 58.0 ± 4.4 $^{a}$ |
| | $^{114}$Cd _E | 4.76 ± 0.33 $^{b}$ | 2.39 ± 0.33 $^{a}$ | 7.15 ± 0.58 $^{a}$ | 52.2 ± 4.9 $^{a}$ | 7.6 ± 1.1 $^{a}$ | 59.8 ± 5.1 $^{a}$ |
| $^{114}$Cd | - | 5.54 ± 0.63 $^{ab}$ | 1.61 ± 0.11 $^{a}$ | 7.15 ± 0.74 $^{a}$ | 47.6 ± 2.0 $^{a}$ | 11.4 ± 4.8 $^{a}$ | 59.1 ± 5.6 $^{a}$ |
| | $^{111}$Cd | 5.38 ± 0.06 $^{a}$ | 1.90 ± 0.28 $^{ab}$ | 7.28 ± 0.33 $^{a}$ | 47.9 ± 2.4 $^{a}$ | 8.9 ± 5.0 $^{a}$ | 56.8 ± 4.5 $^{a}$ |
| | $^{111}$Cd _E | 4.89 ± 0.19 $^{b}$ | 2.32 ± 0.11 $^{b}$ | 7.21 ± 0.27 $^{a}$ | 48.7 ± 2.2 $^{a}$ | 21.8 ±14.6 $^{a}$ | 70.4 ±13.5 $^{a}$ |
| 1$^{st}$ Cd | - | 5.74 ± 0.35 $^{a}$ | 1.66 ± 0.13 $^{a}$ | 7.31 ± 0.50 $^{a}$ | 53.1 ± 3.4 $^{a}$ | 15.5 ± 5.2 $^{a}$ | 68.7 ± 8.1 $^{a}$ |
| (Σ) | 2$^{nd}$ Cd | 5.62 ± 0.11 $^{a}$ | 1.74 ± 0.16 $^{a}$ | 7.36 ± 0.18 $^{a}$ | 50.5 ± 1.9 $^{a}$ | 6.9 ± 2.6 $^{a}$ | 57.4 ± 2.9 $^{a}$ |
| | 2$^{nd}$ Cd_E | 4.83 ± 0.18 $^{b}$ | 2.35 ± 0.16 $^{b}$ | 7.18 ± 0.30 $^{a}$ | 50.4 ± 2.6 $^{a}$ | 14.7 ± 7.3 $^{a}$ | 65.1 ± 7.0 $^{a}$ |
| | | Mg | | | K in Stroma+ | | |
| | | | | | All data | ^ 1$^{st}$ ^ | All data $^{§}$ |
| $^{111}$Cd | - | 1415 ±112 $^{a}$ | 15.1 ± 4.7 $^{a}$ | 1430 ±116 $^{a}$ | 33.5 ± 4.5 $^{a}$ | 30.6 ± 4.7 $^{a}$ | 1 ± 0 $^{a}$ |
| | - ^ | | 10.5 ± 2.0 $^{a}$ | | | | |
| | $^{114}$Cd | 1358 ± 30 $^{a}$ | 11.2 ± 1.5 $^{a}$ | 1369 ± 31 $^{a}$ | 22.8 ± 7.0 $^{a}$ | 15.9 ± 1.2 $^{b}$ | 0.66 ± 0.13 $^{b}$ |
| | $^{114}$Cd _E | 1251 ± 51 $^{a}$ | 9.9 ± 1.0 $^{a}$ | 1261 ± 51 $^{a}$ | 22.2 ± 5.9 $^{a}$ | 16.4 ± 1.1 $^{b}$ | 0.65 ± 0.11 $^{b}$ |
| $^{114}$Cd | - | 1276 ± 47 $^{a}$ | 19.0 ± 3.6 $^{a}$ | 1295 ± 47 $^{a}$ | 24.3 ± 4.4 $^{a}$ | 19.9 ± 0.9 $^{a}$ | 1 ± 0 $^{a}$ |
| | - ^ | | 15.5 ± 1.2 $^{a}$ | | | | |
| | $^{111}$Cd | 1174 ± 51 $^{a}$ | 14.9 ± 1.4 $^{a}$ | 1189 ± 52 $^{a}$ | 19.5 ± 3.2 $^{a}$ | 16.7 ± 2.2 $^{ab}$ | 0.82 ± 0.09 $^{ab}$ |
| | $^{111}$Cd _E | 1157 ± 20 $^{a}$ | 14.7 ± 1.2 $^{a}$ | 1171 ± 21 $^{a2}$ | 20.0 ± 4.2 $^{a}$ | 15.8 ± 0.8 $^{b}$ | 0.82 ± 0.05 $^{b}$ |
| 1$^{st}$ Cd | - | 1345 ± 62 $^{a}$ | 17.0 ± 2.8 $^{a}$ | 1362 ± 63 $^{a}$ | 28.9 ± 3.4 $^{a}$ | 25.3 ± 3.2 $^{a}$ | 1 ± 0 $^{a}$ |
| (Σ) | - ^ | | 13.0 ± 1.5 $^{a}$ | | | | |
| | 2$^{nd}$ Cd | 1266 ± 44 $^{a}$ | 13.0 ± 1.2 $^{a}$ | 1279 ± 44 $^{a}$ | 21.1 ± 3.6 $^{a}$ | 16.3 ± 1.1 $^{b}$ | 0.74 ± 0.08 $^{b}$ |
| | 2$^{nd}$ Cd_E | 1204 ± 31 $^{a}$ | 12.3 ± 1.2 $^{a}$ | 1216 ± 31 $^{a}$ | 21.1 ± 3.4 $^{a}$ | 16.1 ± 0.6 $^{b}$ | 0.74 ± 0.06 $^{b}$ |

Barley plants were grown on initial Cd ($^{111}$Cd or $^{114}$Cd, 80 µM); at this stage, plants and chloroplasts accumulated Cd and other metals *in vivo*. Chloroplasts were isolated from the first (major) and second (minor) leaves. Chloroplasts incubated *in vitro* for 1.5 h in a buffer with late Cd ($^{114}$Cd or $^{111}$Cd, 25 µM). After incubation, chloroplast were washed with a same buffer; in one of variants, EDTA was added to washing buffer. Σ – combined data of both reciprocal experiments: 1$^{st}$ Cd – initial Cd, 2$^{nd}$ Cd – late Cd. *In vitro* variants: "-" – no Cd in incubation buffer, no EDTA in washing buffer; "Cd" – incubation with the corresponding Cd isotope, no EDTA in washing buffer "Cd_E" – incubation with the corresponding Cd isotope and post-washing with EDTA. Cd is shown as sum of the isotopes ($^{111}$Cd + $^{114}$Cd). Stroma+ and $^{\#}$ – see Table S7.

^ – excluding single data point(s). A couple of atypical results (1$^{st}$ exp. $^{111}$Cd/ $^{114}$Cd and 3$^{rd}$ exp. $^{114}$Cd/ $^{111}$Cd) insignificantly increased Mg content in control stroma; without these atypical values, Mg contents were very similar in all variants of stroma.

to Table S9 (continued):
K content determined in stroma solely while represented in three ways. 1st repeat of the experiment slightly differed from the rest repeats: amount of chloroplasts was smaller and they were introduced to ice-cold incubation buffer (typically, pre-warmed to the incubation temperature); therefore, some values of 1st repeat differed from values in 2-4th repeats. In 1st repeat, K contents demonstrated same tendency and larger absolute values making differences insignificant. Significant difference demonstrated in two ways. ^ 1st ^ – excluding data of 1st repeats (repeats 2-4 calculated). § – in each repeat of the experiment, control variant ("-") accepted as 1 and other three variants calculated as portion of 1.
a-c – significant difference between variants of *in vitro* treatment, $p < 0.05$. Close to significant level: 1 – p=0.056; 2 – p=0.052. Differences between reciprocal variants were insignificant.
Means ± SE.

Table S10. Changes of Mn amd some other metals in barley chloroplasts during Cd accumulation *in vitro*. Unpublished data from previous experiment described in (Lysenko et al. 2019 https://doi.org/10.1007/s11120-018-0528-6 )

| | Metal contents. nmol/mg Chl (#) | | | |
|---|---|---|---|---|
| | *In vivo* | *In vitro* | | |
| Cd. µM : | - | 5 | 100 | 100 |
| Washing buffer: | | | | 2 mM EDTA |
| Mn | | | | |
| Thylakoids | 9.31 | 2.24 | 0.96 | 0.00 |
| | 9.17 | 3.34 | 1.68 | 0.00 |
| | a | b | bc | c |
| Stroma+ | 2.12 | 1.65 | 1.84 | 2.72 |
| | 2.34 | 2.17 | 1.84 | 2.54 |
| | ab | ab | a | b |
| Zn | | | | |
| Thylakoids | 1.72 | 1.98 | 1.59 | 0.46 |
| | 1.35 | 1.27 | 0.94 | 0.37 |
| | a | ab | ab | b |
| Stroma+ | 1.93 | 1.08 | 0.50 | 4.76 |
| | 1.11 | 1.60 | 2.58 | 4.26 |
| | a | a | ab | b |
| Cu | | | | |
| Thylakoids | 2.10 | 0.60 | 6.77 | 0.81 |
| | 2.78 | 1.57 | 1.73 | 0.16 |
| | a | a | a | a |
| Stroma+ | 0.67 | 0.61 | 1.00 | 1.19 |
| | 0.39 | 1.19 | 1.29 | 2.13 |
| | a | a | a | a |

The previous experiment was very similar to the current experiment (Table S9, Fig. 5) with two substantial differences. In the previous experiment:

- *In vivo*, barley plants were grown with no Cd addition.
- *In vitro*, Cd accumulation by chloroplasts performed in Tris-based buffer.

Designations are the same as in Table S9. a-c – significant differences.
