## Supplemental Tables 6_7 complete for "Common excluder barley has more than one mechanism to remove Cd from chloroplasts"

**Source:** bioRxiv

Table S6 (complete). Changes of metal contents in organs of barley plants during eight days at Cd-containing mineral medium.

| Me | Organ / Day | nmol/ g FW | | | nmol/ g DW | | | nmol/ organ | | |
|---|---|---|---|---|---|---|---|---|---|---|
| | | $^{111}$Cd/ $^{114}$Cd | $^{114}$Cd/ $^{111}$Cd | Σ | $^{111}$Cd/ $^{114}$Cd | $^{114}$Cd/ $^{111}$Cd | Σ | $^{111}$Cd/ $^{114}$Cd | $^{114}$Cd/ $^{111}$Cd | Σ |
| **Cu** | Roots | | | | | | | | | |
| | 8 | 12,4 ± 2,0 a | 13,6 ± 1,7 a | 13,0 ± 1,2 a | 100 ± 12 a | 108 ± 11ab | 104 ± 8ab | 0,517 ± 0,055 a | 0,585 ± 0,074 a | 0,551 ± 0,045 a |
| | 12 | 12,4 ± 0,8 a | 13,6 ± 0,9 a | 13,0 ± 0,6 a | 99 ± 6 a | 110 ± 5 a | 105 ± 4 a | 0,710 ± 0,023 b | 0,813 ± 0,042 b | 0,762 ± 0,028 b |
| | 16 | 11,2 ± 0,4 a | 12,1 ± 0,9 a | 11,6 ± 0,5 a | 87 ± 3 a | 92 ± 5 b | 90 ± 3 b | 0,773 ± 0,050 b | 0,751 ± 0,031ab | 0,762 ± 0,028 b |
| | Stem+ | | | | | | | | | |
| | 8 | 7,88 ± 0,36 a | 6,89 ± 0,99 a | 7,39 ± 0,52 a | 71,0 ± 2,7 a | 62,4 ± 9,8 a | 66,7 ± 5,0 a | 0,461 ± 0,023 a | 0,381 ± 0,060 a | 0,421 ± 0,033 a |
| | 12 | 3,97 ± 0,65 b | 3,91 ± 0,71 b | 3,94 ± 0,45 b | 34,0 ± 5,6 b | 33,6 ± 5,9 b | 33,8 ± 3,8 b | 0,537 ± 0,072 a | 0,607 ± 0,137ab | 0,572 ± 0,074ab |
| | 16 | 3,83 ± 0,16 b | 3,38 ± 0,66 b | 3,61 ± 0,33 b | 29,3 ± 1,8 b | 26,3 ± 4,7 b | 27,8 ± 2,4 b | 0,767 ± 0,065 b | 0,682 ± 0,115 b | 0,725 ± 0.064 b |
| | 1st leaf | | | | | | | | | |
| | 8 | 3,61 ± 0,66 a | 3,50 ± 0,70 a | 3,56 ± 0,46 a | 33,9 ± 6,5 a | 32,4 ± 6,5 a | 33,2 ± 4,3 a | 0,369 ± 0.065 a | 0,351 ± 0,069 a | 0,360 ± 0,045 a |
| | 12 | 3,20 ± 0,67 a | 2,68 ± 0,16 a | 2,94 ± 0,34 a | 27,5 ± 5,9 a | 23,3 ± 1,3 a | 25,4 ± 2,9 a | 0,324 ± 0,072 a | 0,284 ± 0,012 a | 0,304 ± 0,035ab |
| | 16 | 2,71 ± 0,55 a | 2,60 ± 0,28 a | 2,65 ± 0,29 a | 22,2 ± 4,5 a | 22,5 ± 2,0 a | 22,4 ± 2,3 a | 0,235 ± 0,042 a | 0,256 ± 0,028 a | 0,245 ± 0,024 b |
| **Mn** | Roots | | | | | | | | | |
| | 8 | 41,2 ± 3,0 a | 51,2 ± 3,2 a | 46,2 ± 2,7 a | 336 ± 16 a | 410 ± 16 a | 373 ± 16 a | 1,78 ± 0,15 a | 2,21 ± 0,15 a | 1,99 ± 0,12 a |
| | 12 | 36,2 ± 1,2 a | 34,7 ± 1,8 b | 35,5 ± 1,0 b | 290 ± 9 b | 284 ± 20 b | 287 ± 11 b | 2,11 ± 0,17 a | 2,12 ± 0,21 a | 2,11 ± 0,13 a |
| | 16 | 41,0 ± 2,4 a | 37,5 ± 3,2 b | 39,3 ± 2,0ab | 319 ± 7 a | 284 ± 14 b | 302 ± 9 b | 2,81 ± 0,12 b | 2,38 ± 0,29 a | 2,60 ± 0,16 b |
| | Stem+ | | | | | | | | | |
| | 8 | 50,9 ± 1,7 a | 48,6 ± 1,3 a | 49,7 ± 1,1 a | 462 ± 34 a | 437 ± 19 a | 450 ± 19 a | 2,99 ± 0,21 a | 2,66 ± 0,11 a | 2,83 ± 0,12 a |
| | 12 | 32,9 ± 0,8 b | 35,3 ± 1,2 c | 34,1 ± 0,8 b | 282 ± 7 b | 306 ± 14 c | 294 ± 8 c | 4,58 ± 0,22 b | 5,20 ± 0,19 b | 4,89 ± 0,17 b |
| | 16 | 40,0 ± 0,9 c | 39,5 ± 1,5 c | 39,7 ± 0,8 c | 305 ± 7 c | 314 ± 16 c | 309 ± 8 c | 7,96 ± 0,43 c | 8,25 ± 0,64 c | 8,11 ± 0,37 c |
| | 1st leaf | | | | | | | | | |
| | 8 | 58,7 ± 2,7 a | 60,6 ± 1,6 a | 59,6 ± 1,5 a | 548 ± 19 a | 561 ± 9 a | 554 ± 10 a | 6,03 ± 0,27 a | 6,10 ± 0,14 a | 6,07 ± 0,15 a |
| | 12 | 71,3 ± 2,1 b | 69,6 ± 4,1ac | 70,4 ± 2,2 b | 609 ± 23 a | 601 ± 24 a | 605 ± 16 b | 7,21 ± 0,43 b | 7,37 ± 0,29 b | 7,29 ± 0,24 b |
| | 16 | 87,4 ± 2,4 c | 79,9 ± 3,8 c | 83,6 ± 2,5 c | 718 ± 16 c | 695 ± 16 c | 706 ± 12 c | 7,67 ± 0,18 b | 7,86 ± 0,37 b | 7,77 ± 0,20 b |

Table S6 (complete, continued). Changes of metal contents in organs of barley plants during eight days at Cd-containing mineral medium.

| Me | Organ / Day | µmol/ g FW | | | µmol/ g DW | | | nmol/ organ | | |
|---|---|---|---|---|---|---|---|---|---|---|
| | | $^{111}$Cd/ $^{114}$Cd | $^{114}$Cd/ $^{111}$Cd | Σ | $^{111}$Cd/ $^{114}$Cd | $^{114}$Cd/ $^{111}$Cd | Σ | $^{111}$Cd/ $^{114}$Cd | $^{114}$Cd/ $^{111}$Cd | Σ |
| **Zn** | Roots | | | | | | | | | |
| | 8 | 1,89 ± 0,45 a | 1,34 ± 0,21 a | 1,62 ± 0,25 a | 16,2 ± 4,5ac | 10,9 ± 1,9 a | 13,6 ± 2,5 a | 85 ± 25 a | 58 ± 9 a | 72 ± 13 a |
| | 12 | 2,26 ± 0,24 a | 2,27 ± 0,09 b | 2,27 ± 0,12 b | 18,3 ± 2,4 a | 18,6 ± 1,2 b | 18,5 ± 1,2 a | 131 ± 15 a | 139 ± 13 b | 135 ± 9 b |
| | 16 | 3,47 ± 0,26 c | 3,36 ± 0,31 c | 3,42 ± 0,19 c | 27,4 ± 2,8 c | 25,5 ± 1,6 c | 26,5 ± 1,6 c | 247 ± 34 c | 214 ± 29 c | 230 ± 22 c |
| | Stem+ | | | | | | | | | |
| | 8 | 0,239 ± 0,055 a | 0,172 ± 0,030 a | 0,205 ± 0,031 a | 2,24 ± 0,61 a | 1,56 ± 0,29 a | 1,90 ± 0,34 a | 14,3 ± 3,8 a | 9,5 ± 1,8 a | 11,9 ± 2,1 a |
| | 12 | 0,350 ± 0,024 a | 0,326 ± 0,021 b | 0,338 ± 0,016 b | 3,00 ± 0,22 a | 2,81 ± 0,15 b | 2,91 ± 0,13 b | 49,2 ± 5,5 b | 49,1 ± 6,5 b | 49,2 ± 4,0 b |
| | 16 | 0,595 ± 0,072 c | 0,661 ± 0,117 c | 0,628 ± 0,066 c | 4,54 ± 0,55 c | 5,17 ± 0,78 c | 4,86 ± 0,46 c | 120,4 ± 18,4 c | 136,6 ± 21,4 c | 128,5 ± 13,6 c |
| | 1$^{st}$ leaf | | | | | | | | | |
| | 8 | 0,171 ± 0,052 a | 0,112 ± 0,020 a | 0,142 ± 0,028 a | 1,66 ± 0,55 a | 1,05 ± 0,21 a | 1,35 ± 0,30 a | 17,8 ± 5,7 a | 11,4 ± 2,3 a | 14,6 ± 3,1 a |
| | 12 | 0,176 ± 0,027 a | 0,163 ± 0,013 a | 0,170 ± 0,014 a | 1,51 ± 0,25 a | 1,42 ± 0,13 a | 1,47 ± 0,13 a | 17,6 ± 2,5 a | 17,6 ± 1,9ab | 17,6 ± 1,5 a |
| | 16 | 0,224 ± 0,027 a | 0,255 ± 0,036 b | 0,239 ± 0,022 b | 1,85 ± 0,23 a | 2,21 ± 0,29 b | 2,03 ± 0,19 b | 19,7 ± 2,3 a | 25,3 ± 3,9 b | 22,5 ± 2,4 a |
| **Fe** | Roots | | | | | | | | | |
| | 8 | 3,95 ± 0,81 a | 4,45 ± 0,57 a | 4,20 ± 0,47 a | 32,6 ± 6,8 a | 35,6 ± 4,5 a | 34,1 ± 3,9 a | 172 ± 36 a | 193 ± 26 a | 182 ± 21 a |
| | 12 | 11,39 ± 1,63 b | 8,96 ± 1,06 b | 10,17 ± 1,00 b | 91,9 ± 14,2 c | 72,4 ± 7,2 c | 82,1 ± 8,2 b | 658 ± 92 b | 539 ± 66 b | 599 ± 57 b |
| | 16 | 15,43 ± 0,63 c | 12,41 ± 2,01 c | 13,92 ± 1,11 c | 121,4 ± 8,7 c | 94,8 ± 14,7 c | 108,1 ± 9,2 c | 1085 ± 118 c | 813 ± 183 c | 949 ± 112 c |
| | Stem+ | | | | | | | | | |
| | 8 | 0,114 ± 0,012 a | 0,124 ± 0,005 a | 0,119 ± 0,006 a | 1,02 ± 0,08 a | 1,11 ± 0,04 a | 1,07 ± 0,05 a | 6,6 ± 0,5 a | 6,8 ± 0,3 a | 6,7 ± 0,3 a |
| | 12 | 0,165 ± 0,005 b | 0,173 ± 0,023ac | 0,169 ± 0,011 c | 1,41 ± 0,03 b | 1,50 ± 0,22ab | 1,46 ± 0,11 b | 23,0 ± 1,5 b | 25,2 ± 2,6 b | 24,1 ± 1,5 b |
| | 16 | 0,198 ± 0,005 c | 0,178 ± 0,006 c | 0,188 ± 0,005 c | 1,50 ± 0,03 b | 1,41 ± 0,03 b | 1,46 ± 0,02 b | 39,5 ± 2,7 c | 37,1 ± 2,5 c | 38,3 ± 1,8 c |
| | 1$^{st}$ leaf | | | | | | | | | |
| | 8 | 0,149 ± 0,015 a | 0,178 ± 0,008 a | 0,164 ± 0,009 a | 1,38 ± 0,10 a | 1,64 ± 0,04 a | 1,51 ± 0,07 a | 15,3 ± 1,4 a | 17,9 ± 0,6 a | 16,6 ± 0,8 a |
| | 12 | 0,234 ± 0,043ab | 0,219 ± 0,010 b | 0,226 ± 0,021 b | 2,00 ± 0,37ab | 1,90 ± 0,10 b | 1,95 ± 0,18 b | 24,1 ± 5,4 a | 23,3 ± 1,4 b | 23,7 ± 2,6 b |
| | 16 | 0,219 ± 0,012 b | 0,258 ± 0,039ab | 0,239 ± 0,020 b | 1,81 ± 0,11 b | 2,22 ± 0,28ab | 2,01 ± 0,16 b | 19,4 ± 1,4 a | 25,5 ± 4,2ab | 22,4 ± 2,3 b |

Table S6 (complete, continued). Changes of metal contents in organs of barley plants during eight days at Cd-containing mineral medium.

| Me | Organ / Day | µmol/ g FW | | | µmol/ g DW | | | nmol/ organ | | |
|---|---|---|---|---|---|---|---|---|---|---|
| | | $^{111}Cd/ ^{114}Cd$ | $^{114}Cd/ ^{111}Cd$ | Σ | $^{111}Cd/ ^{114}Cd$ | $^{114}Cd/ ^{111}Cd$ | Σ | $^{111}Cd/ ^{114}Cd$ | $^{114}Cd/ ^{111}Cd$ | Σ |
| **Mg** | Roots | | | | | | | | | |
| | 8 | 7,72 ± 0,26 a | 7,39 ± 0,26 a | 7,56 ± 0,18 a | 63,6 ± 3,4 a | 59,4 ± 2,1 a | 61,5 ± 2,0 a | 335 ± 26 a | 318 ± 8 a | 327 ± 13 a |
| | 12 | 6,81 ± 0,27 b | 6,55 ± 0,40 a | 6,68 ± 0,23 b | 54,4 ± 1,2 b | 53,2 ± 2,6 a | 53,8 ± 1,4 b | 394 ± 25ab | 394 ± 32 b | 394 ± 19 b |
| | 16 | 6,66 ± 0,50ab | 6,98 ± 0,71 a | 6,82 ± 0,41ab | 51,7 ± 2,0 b | 52,8 ± 2,8 a | 52,2 ± 1,6 b | 456 ± 25 b | 434 ± 35 b | 445 ± 21 b |
| | Stem+ | | | | | | | | | |
| | 8 | 8,70 ± 0,20 a | 8,56 ± 0,35ab | 8,63 ± 0,19 a | 78,5 ± 2,0 a | 76,7 ± 2,3 a | 77,6 ± 1,5 a | 512 ± 34 a | 466 ± 10 a | 489 ± 19 a |
| | 12 | 8,42 ± 0,19 a | 7,63 ± 0,52 a | 8,03 ± 0,29 a | 72,1 ± 1,4 b | 66,2 ± 5,3 a | 69,2 ± 2,8 b | 1172 ± 61 b | 1117 ± 46 b | 1144 ± 37 b |
| | 16 | 9,69 ± 0,18 b | 8,95 ± 0,23 b | 9,32 ± 0,19 b | 73,9 ± 1,8ab | 71,1 ± 2,8 a | 72,5 ± 1,6 b | 1936 ± 132 c | 1871 ± 142 c | 1904 ± 92 c |
| | $1^{st}$ leaf | | | | | | | | | |
| | 8 | 6,86 ± 0,21 a | 7,01 ± 0,21 a | 6,93 ± 0,14 a | 64,1 ± 1,3ab | 65,0 ± 2,1 a | 64,5 ± 1,2 a | 705 ± 21 a | 705 ± 22 a | 705 ± 14 a |
| | 12 | 7,22 ± 0,14 a | 6,51 ± 0,40 a | 6,86 ± 0,23 a | 61,7 ± 1,9 a | 56,4 ± 3,3 a | 59,0 ± 2,0 c | 729 ± 36 a | 695 ± 50 a | 712 ± 30 a |
| | 16 | 8,58 ± 0,34 b | 7,41 ± 0,29 a | 8,00 ± 0,28 b | 70,5 ± 2,7 b | 64,5 ± 1,7 a | 67,5 ± 1,8 a | 753 ± 24 a | 728 ± 24 a | 741 ± 16 a |
| **Ca** | Roots | | | | | | | | | |
| | 8 | 8,09 ± 0,34 a | 7,58 ± 0,27 a | 7,84 ± 0,22 a | 66,5 ± 2,3 a | 60,8 ± 1,4 a | 63,6 ± 1,6 a | 349 ± 17 a | 327 ± 14 a | 338 ± 11 a |
| | 12 | 9,08 ± 0,49 a | 9,87 ± 0,78 c | 9,48 ± 0,45 b | 72,8 ± 4,1 a | 80,8 ± 7,9 c | 76,8 ± 4,4 c | 531 ± 54 b | 609 ± 92 c | 570 ± 52 b |
| | 16 | 11,06 ± 0,20 c | 10,87 ± 0,77 c | 10,96 ± 0,38 c | 86,5 ± 2,7 c | 82,7 ± 3,3 c | 84,6 ± 2,1 c | 771 ± 62 c | 690 ± 81 c | 731 ± 50 c |
| | Stem+ | | | | | | | | | |
| | 8 | 9,20 ± 0,45 a | 8,37 ± 0,36 a | 8,78 ± 0,31 a | 83 ± 6 a | 75 ± 3 a | 79 ± 3 a | 545 ± 53 a | 457 ± 22 a | 501 ± 31 a |
| | 12 | 9,12 ± 1,10 a | 8,46 ± 0,27 a | 8,79 ± 0,55 a | 78 ± 10 a | 73 ± 2 a | 76 ± 5 a | 1249 ± 128 b | 1262 ± 110 b | 1256 ± 80 b |
| | 16 | 15,43 ± 1,98 b | 12,30 ± 0,66 b | 13,86 ± 1,12 b | 117 ± 13 b | 98 ± 6 b | 107 ± 8 b | 3113 ± 486 c | 2581 ± 247 c | 2847 ± 272 c |
| | $1^{st}$ leaf | | | | | | | | | |
| | 8 | 14,7 ± 1,2 a | 13,6 ± 0,7 a | 14,2 ± 0,7 a | 138 ± 11 a | 126 ± 6 a | 132 ± 6 a | 1512 ± 117 a | 1366 ± 69 a | 1439 ± 68 a |
| | 12 | 21,0 ± 1,4 b | 20,1 ± 1,6 b | 20,5 ± 1,0 b | 179 ± 11 b | 173 ± 10 b | 176 ± 7 b | 2121 ± 161 c | 2121 ± 125 c | 2121 ± 96 b |
| | 16 | 28,9 ± 1.6 c | 25,7 ± 0,4 c | 27,3 ± 0,9 c | 238 ± 14 c | 224 ± 7 c | 231 ± 8 c | 2537 ± 137 c | 2543 ± 148 c | 2540 ± 95 c |

Table S6 (complete, continued). Changes of metal contents in organs of barley plants during eight days at Cd-containing mineral medium.

| Me | Organ / Day | µmol/ g FW | | | µmol/ g DW | | | µmol/ organ | | |
|---|---|---|---|---|---|---|---|---|---|---|
| | | $^{111}$Cd/ $^{114}$Cd | $^{114}$Cd/ $^{111}$Cd | Σ | $^{111}$Cd/ $^{114}$Cd | $^{114}$Cd/ $^{111}$Cd | Σ | $^{111}$Cd/ $^{114}$Cd | $^{114}$Cd/ $^{111}$Cd | Σ |
| **Na** | Roots | | | | | | | | | |
| | 8 | 47,0 ± 4,8 a | 42,5 ± 3,2 a | 44,5 ± 2,7 a | 372 ± 34 a | 342 ± 28 a | 355 ± 21 a | 1,90 ± 0,10 a | 1,82 ± 0,11 a | 1,86 ± 0,07 a |
| | 12 | 35,3 ± 2,1 a | 33,1 ± 2,9 a | 34,2 ± 1,7 b | 282 ± 14 b | 268 ± 16ab[1] | 275 ± 10 b | 2,06 ± 0,20ab | 1,96 ± 0,07 a | 2,01 ± 0,10ab |
| | 16 | 35,6 ± 2,4 a | 34,7 ± 3,5 a | 35,1 ± 2,0 b | 278 ± 16 b | 263 ± 16 b | 270 ± 11 b | 2,46 ± 0,19 b | 2,15 ± 0,13 a | 2,30 ± 0,12 b |
| | Stem+ | | | | | | | | | |
| | 8 | 12,7 ± 1,4ab | 10,3 ± 1,0 a | 11,5 ± 0,9ab | 115 ± 13ab | 93 ± 11 a | 104 ± 9ab | 0,76 ± 0,13 a | 0,56 ± 0,06 a | 0,66 ± 0,07 a |
| | 12 | 10,3 ± 0,9 a | 10,0 ± 1,0 a | 10,2 ± 0,6 a | 88 ± 7 a | 87 ± 10 a | 88 ± 6 a | 1,43 ± 0,10 b | 1,45 ± 0,07 b | 1,44 ± 0,06 b |
| | 16 | 15,9 ± 1,0 b | 12,3 ± 0,6 a | 14,1 ± 0,8 b | 121 ± 8 b | 98 ± 5 a | 110 ± 6 b | 3,17 ± 0,27 c | 2,58 ± 0,22 c | 2,87 ± 0,19 c |
| | 1st leaf | | | | | | | | | |
| | 8 | 14,9 ± 1,7 a | 12,7 ± 1,1 a | 13,8 ± 1,0 a | 140 ± 16 a | 118 ± 12 a | 129 ± 10 a | 1,53 ± 0,17 a | 1,29 ± 0,13 a | 1,41 ± 0,11 a |
| | 12 | 17,4 ± 0,8 a | 17,0 ± 1,1 c | 17,2 ± 0,6 b | 149 ± 9 a | 147 ± 10ac | 148 ± 6 a | 1,75 ± 0,09 a | 1,80 ± 0,09 c | 1,78 ± 0,06 b |
| | 16 | 27,2 ± 1,0 c | 19,8 ± 1,2 c | 23,5 ± 1,4 c | 224 ± 8 c | 173 ± 9 c | 198 ± 10 c | 2,40 ± 0,10 c | 1,96 ± 0,15 c | 2,18 ± 0,11 c |
| **K** | Roots | | | | | | | | | |
| | 8 | 88 ± 8 a | 103 ± 2 a | 95 ± 5 a | 724 ± 64 a | 823 ± 11 a | 774 ± 35 a | 3,78 ± 0,31 a | 4,42 ± 0,11 a | 4,10 ± 0,19 a |
| | 12 | 94 ± 9ab | 103 ± 5 a | 99 ± 5ab | 754 ± 70 a | 840 ± 44 a | 797 ± 42 a | 5,47 ± 0,58 b | 6,24 ± 0,52 c | 5,86 ± 0,39 b |
| | 16 | 111 ± 2 b | 107 ± 6 a | 109 ± 3 b | 865 ± 22 a | 817 ± 29 a | 841 ± 19 a | 7,68 ± 0,54 c | 6,79 ± 0,69 c | 7,24 ± 0,44 c |
| | Stem+ | | | | | | | | | |
| | 8 | 146 ± 2 a | 138 ± 5 a | 142 ± 3 a | 1322 ± 54 a | 1235 ± 35 a | 1278 ± 34 a | 8,58 ± 0,43 a | 7,50 ± 0,10 a | 8,04 ± 0,27 a |
| | 12 | 209 ± 4 b | 187 ± 9 b | 198 ± 6 b | 1797 ± 65 c | 1610 ± 68 c | 1704 ± 54 c | 29,17 ± 1,69 b | 28,01 ± 2,98 b | 28,59 ± 1,63 b |
| | 16 | 233 ± 3 c | 239 ± 12 c | 236 ± 6 c | 1783 ± 63 c | 1906 ± 142 c | 1844 ± 76 c | 46,81 ± 3,63 c | 50,40 ± 5,46 c | 48,60 ± 3,15 c |
| | 1st leaf | | | | | | | | | |
| | 8 | 171 ± 12 a | 183 ± 5 a | 177 ± 6 a | 1610 ± 142 a | 1700 ± 52 a | 1655 ± 73 a | 17,61 ± 1,30 a | 18,48 ± 0,69 a | 18,05 ± 0,71 a |
| | 12 | 208 ± 6 c | 230 ± 8 c | 219 ± 6 c | 1773 ± 50 a | 1999 ± 79 c | 1886 ± 58 c | 21,04 ± 1,33 a | 24,59 ± 1,40 b | 22,82 ± 1,08 b |
| | 16 | 179 ± 28ac | 208 ± 5 b | 193 ± 14ac | 1468 ± 226 a | 1814 ± 38ac | 1641 ± 123ac | 15,80 ± 2,54 a | 20,55 ± 1,02 a | 18,17 ± 1,51 a |

The experiment is described in Table S4. Me – metal. Roots – of a single plant (per organ), stem+ – stem with leaf sheaths, 1st leaf - first leaf, leaf blade. SI prefixes colored to ease perception. Reciprocal experiments: $^{111}$Cd/ $^{114}$Cd and $^{114}$Cd/ $^{111}$Cd; Σ – combined data of both reciprocal experiments: initial Cd/ late Cd. a-c – significant differences between plants of diverse ages in a single variant, $p < 0.05$; 1 – difference significant at $p=0.0517$. No significant differences observed between reciprocal variants. Means ± SE

Table S7 (complete). Changes of metal contents in chloroplasts of barley first leaves during eight days at Cd-containing mineral medium.

| Me/ variant | Day | Fractions of chloroplasts | | | | Chloroplasts | |
|---|---|---|---|---|---|---|---|
| | | Thylakoids | | Stroma + envelope | | nmol/mg Chl | % from leaves [×] |
| | | nmol/mg Chl | % [&] | nmol/mg Chl [#] | % [&] | | |
| Cd | | | | | | | |
| 111/ 114 | 8 | 1.13 ± 0.04 [a] | 83.3 ± 1.3 | 0.23 ± 0.02 | 16.7 ± 1.3 | 1.36 ± 0.05 [a] | 2.16 ± 0.10 [a] |
| | 16 | 1.27 ± 0.04 [b] | 84.9 ± 1.1 | 0.23 ± 0.02 | 15.1 ± 1.1 | 1.50 ± 0.06 [a] | 1.08 ± 0.04 [b] |
| | 16^ | 0.96 ± 0.06 [c] | | 0.17 ± 0.01 [1] | | 1.13 ± 0.06 [b] | |
| 114/ 111 | 8 | 1.13 ± 0.01 [a] | 82.7 ± 0.9 | 0.24 ± 0.02 | 17.3 ± 0.9 | 1.36 ± 0.02 [ab] | 2.01 ± 0.11 [a] |
| | 16 | 1.28 ± 0.06 [b] | 84.3 ± 0.7 | 0.24 ± 0.02 | 15.7 ± 0.7 | 1.52 ± 0.07 [a] | 1.10 ± 0.08 [b] |
| | 16^ | 1.07 ± 0.06 [a] | | 0.20 ± 0.01 [2] | | 1.26 ± 0.07 [b] | |
| Σ | 8 | 1.13 ± 0.02 [a] | 83.0 ± 0.8 | 0.23 ± 0.01 [a] | 17.0 ± 0.8 | 1.36 ± 0.03 [a] | 2.08 ± 0.07 [a] |
| | 16 | 1.28 ± 0.03 [b] | 84.6 ± 0.6 | 0.23 ± 0.01 [a] | 15.4 ± 0.6 | 1.51 ± 0.05 [b] | 1.09 ± 0.04 [b] |
| | 16^ | 1.01 ± 0.04 [c] | | 0.18 ± 0.01 [b] | | 1.20 ± 0.05 [c] | |
| Cu | | | | | | | |
| 111/ 114 | 8 | 1.45 ± 0.10 [a] | 79.6 ± 2.5 | 0.37 ± 0.05 | 20.4 ± 2.5 | 1.82 ± 0.10 [a] | 87 ± 13 |
| | 16 | 1.08 ± 0.09 [b] | 77.8 ± 3.8 | 0.30 ± 0.05 | 22.2 ± 3.8 | 1.38 ± 0.07 [b] | 68 ± 11 |
| 114/ 111 | 8 | 1.59 ± 0.09 [a] | 80.9 ± 2.8 | 0.39 ± 0.08 | 19.1 ± 2.8 | 1.99 ± 0.15 [a] | 101 ± 29 |
| | 16 | 1.06 ± 0.06 [b] | 71.8 ± 3.2 | 0.42 ± 0.06 | 28.2 ± 3.2 | 1.49 ± 0.09 [b] | 69 ± 9 |
| Σ | 8 | 1.52 ± 0.07 [a] | 80.2 ± 1.8 | 0.38 ± 0.05 | 19.8 ± 1.8 | 1.90 ± 0.09 [a] | 94 ± 15 |
| | 16 | 1.07 ± 0.05 [b] | 74.8 ± 2.6 | 0.36 ± 0.04 | 25.2 ± 2.6 | 1.44 ± 0.05 [b] | 68 ± 7 |
| Mn | | | | | | | |
| 111/ 114 | 8 | 10.1 ± 0.9 | 86.3 ± 1.0 | 1.55 ± 0.03 | 13.7 ± 1.0 | 11.6 ± 0.9 | 30.9 ± 2.6 [a] |
| | 16 | 8.2 ± 0.2 | 84.3 ± 0.8 | 1.54 ± 0.07 | 15.7 ± 0.8 | 9.8 ± 0.2 | 13.2 ± 0.8 [b] |
| 114/ 111 | 8 | 9.5 ± 0.5 | 86.4 ± 0.9 | 1.47 ± 0.04 | 13.6 ± 0.9 | 10.9 ± 0.4 | 25.2 ± 1.8 [a] |
| | 16 | 8.4 ± 0.4 | 84.8 ± 0.9 | 1.50 ± 0.05 | 15.2 ± 0.9 | 9.9 ± 0.3 | 14.3 ± 0.6 [b] |
| Σ | 8 | 9.8 ± 0.5 [a] | 86.4 ± 0.6 [a] | 1.51 ± 0.03 | 13.6 ± 0.6 [a] | 11.3 ± 0.5 [a] | 28.1 ± 1.8 [a] |
| | 16 | 8.3 ± 0.2 [b] | 84.5 ± 0.6 [b] | 1.52 ± 0.04 | 15.5 ± 0.6 [b] | 9.9 ± 0.2 [b] | 13.7 ± 0.5 [b] |
| Fe | | | | | | | |
| 111/ 114 | 8 | 66.6 ± 4.3 | 95.0 ± 1.4 | 3.7 ± 1.2 | 5.0 ± 1.4 | 72.3 ± 5.4 | 78 ± 12 [a] |
| | 16 | 68.5 ± 8.0 | 95.7 ± 0.5 | 3.2 ± 0.6 | 4.3 ± 0.5 | 71.6 ± 8.6 | 38 ± 4 [b] |
| 114/ 111 | 8 | 69.8 ± 3.7 | 95.7 ± 1.8 | 3.3 ± 1.6 | 4.3 ± 1.8 | 73.1 ± 4.3 | 58 ± 6 [a] |
| | 16 | 61.2 ± 2.2 | 94.6 ± 0.5 | 3.5 ± 0.4 | 5.4 ± 0.5 | 64.7 ± 2.3 | 31 ± 4 [b] |
| Σ | 8 | 68.2 ± 2.7 | 95.4 ± 1.1 | 3.5 ± 1.0 | 4.6 ± 1.1 | 72.8 ± 3.1 | 67 ± 7 [a] |
| | 16 | 64.8 ± 4.1 | 95.1 ± 0.4 | 3.3 ± 0.3 | 4.9 ± 0.4 | 68.1 ± 4.3 | 35 ± 3 [b] |
| Mg | | | | | | | |
| 111/ 114 | 8 | 1327 ± 124 | 98.5 ± 0.2 | 19.9 ± 2.5 | 1.5 ± 0.2 | 1347 ± 124 | 30.9 ± 4.2 [a] |
| | 16 | 1281 ± 34 | 98.8 ± 0.3 | 16.2 ± 4.0 | 1.2 ± 0.3 | 1297 ± 37 | 17.8 ± 1.1 [b] |
| 114/ 111 | 8 | 1372 ± 90 | 98.5 ± 0.2 | 20.7 ± 2.2 | 1.5 ± 0.2 | 1393 ± 90 | 27.7 ± 2.2 [a] |
| | 16 | 1281 ± 46 | 98.6 ± 0.3 | 18.3 ± 4.0 | 1.4 ± 0.3 | 1299 ± 49 | 20.4 ± 1.6 [b] |
| Σ | 8 | 1350 ± 73 | 98.5 ± 0.1 | 20.3 ± 1.6 | 1.5 ± 0.1 | 1370 ± 73 | 29.3 ± 2.3 [a] |
| | 16 | 1281 ± 27 | 98.7 ± 0.2 | 17.3 ± 2.7 | 1.3 ± 0.2 | 1298 ± 29 | 19.1 ± 1.0 [b] |
| K | | | | | | | $ |
| 111/ 114 | 8 | n.d. | | 13.5 ± 2.2 | | | 0.013 ± 0.004 |
| | 16 | n.d. | | 49.6 ± 19.4 | | | 0.045 ± 0.023 |
| 114/ 111 | 8 | n.d. | | 9.2 ± 0.8 | | | 0.007 ± 0.001 |
| | 16 | n.d. | | 51.4 ± 21.9 | | | 0.028 ± 0.012 |
| Σ | 8 | n.d. | | 11.4 ± 1.3 [a] | | | 0.010 ± 0.002 |
| | 16 | n.d. | | 50.5 ± 13.8 [b] | | | 0.037 ± 0.013 |

The experiment is described in Table S4; chloroplasts isolated from the first leaves. Cd is shown as the sum of isotopes ($^{111}$Cd + $^{114}$Cd). Me – metal; $^{\&}$ – % in chloroplasts; $^{\#}$ – per mg Chl in chloroplasts from which the stromal fraction was separated; ˟ – see Table S3; $ – portion of K in stroma+envelope (%) of all K in the first leaves; n.d. – not detected. K content in thylakoids is tiny and required more chloroplasts for proper measurement (Lysenko et al. 2019). Reciprocal experiments: 111/114 – $^{111}$Cd/ $^{114}$Cd, 114/111 – $^{114}$Cd/ $^{111}$Cd, Σ – combined data of both reciprocal experiments: initial Cd/ late Cd. ^ - correction for the effect of Chl decrease; for the sum of isotopes, both corrections (Eq. 4 and Eq. 8) resulted in identical values and did not change proportions (%). a-b-c – significant difference between chloroplasts from 8- and 16-day-old plants in a corresponding variant, $p < 0.05$; pairs with no significant difference remained unmarked to ease perception; 1 – difference significant at p=0.0537, 2 – difference significant at p=0.0521. Differences between reciprocal variants were insignificant. Means ± SE.
